# Cryo-EM structure and polar assembly of the PS2 S-layer of *Corynebacterium glutamicum*

**DOI:** 10.1101/2024.09.05.611363

**Authors:** Adrià Sogues, Mike Sleutel, Julienne Petit, Daniela Megrian, Nicolas Bayan, Anne Marie Wehenkel, Han Remaut

**Affiliations:** Structural and Molecular Microbiology, VIB-VUB Center for Structural Biology, VIB, Pleinlaan 2, 1050 Brussels, Belgium; Structural Biology Brussels, Vrije Universiteit Brussel, VUB, Pleinlaan 2, 1050 Brussels, Belgium; Institut Pasteur, Université Paris Cité, CNRS UMR 3528, Bacterial Cell Cycle Mechanisms Unit, F-75015 Paris, France; Institut Pasteur, Université Paris Cité, CNRS UMR 3528, Structural Microbiology Unit, F-75015 Paris, France; Bioinformatics Unit, Institut Pasteur de Montevideo, 11200 Montevideo, Uruguay; Université Paris-Saclay, CEA, CNRS, Institute for Integrative Biology of the Cell (I2BC), Gif-sur-Yvette, France

## Abstract

The polar-growing Corynebacteriales have a complex cell envelope architecture characterized by the presence of a specialized outer membrane composed of mycolic acids. In some Corynebacteriales, this mycomembrane is further supported by a proteinaceous surface layer or ‘S-layer’, whose function, structure and mode of assembly remain largely enigmatic. Here, we isolated *ex vivo* PS2 S-layers from the industrially important *Corynebacterium glutamicum* and determined its atomic structure by 3D cryoEM reconstruction. PS2 monomers consist of a six-helix bundle ‘core’, a three-helix bundle ‘arm’, and a C-terminal transmembrane (TM) helix. The PS2 core oligomerizes into hexameric units anchored in the mycomembrane by a channel-like coiled-coil of the TM helices. The PS2 arms mediate trimeric lattice contacts, crystallizing the hexameric units into an intricate semipermeable lattice. Using pulse-chase live cell imaging, we show that the PS2 lattice is incorporated at the poles, coincident with the actinobacterial elongasome. Finally, phylogenetic analysis shows a paraphyletic distribution and dispersed chromosomal location of PS2 in Corynebacteriales as a result of multiple recombination events and losses. These findings expand our understanding of S-layer biology and enable applications of membrane-supported self-assembling bioengineered materials.

## Introduction

Since the first discovery of a Surface layer (S-layer) over 70 years ago (Houwink, 1953), researchers have identified hundreds of S-layers across nearly every bacterial taxonomic group, and in the majority of Archaea. S-layers are two-dimensional monolayered crystals typically composed of a single (glyco)protein that self-assembles to cover the entire cell surface. Considered one of the most abundant protein families on earth, S-layers are often regulated by strong promoters and have stable mRNA half-lives, accounting for 10 to 30% of total protein synthesis and representing a significant energy cost for the cell (Sleytr, Schuster, Egelseer, & Pum, 2014). Many S-layer proteins share a bipartite architecture comprising a cell envelope binding domain and a crystallization domain that self-assembles into 2D lattices of defined symmetry (Pum, Breitwieser, & Sleytr, 2021). Yet, S-layers often lack discernable sequence or structural homology across different taxonomic groups, indicating their multiple independent emergences across the evolutionary tree of life, likely driven by the advantageous traits they confer as continuous semipermeable non-membraneous layers (Bharat et al., 2020). The reported physiological roles and functions of S-layers are remarkably diverse and may be pleiotropic in many cases (Sleytr et al., 2014),(Beveridge et al., 1997). Research across various organisms suggests S-layers play roles in adhesion (Alp, Kuleaşan, & Korkut Altıntaş, 2020), cell-shape maintenance (C. Zhang et al., 2019), and virulence (Fioravanti et al., 2019), (Assandri, Malamud, Trejo, & Serradell, 2023), or function as molecular sieves (Kügelgen, Cassidy, Dorst, Pagani, & Bharat, 2024) or as a cell envelope supporting exoskeleton (Fioravanti, Mathelie-Guinlet, Dufrêne, Remaut, & Nelson, 2022),(Sogues et al., 2023). S-layer function is often hard to discern. Testament to this is the fact that for numerous characterized S-layers functional data is still lacking as their knockouts show little to no phenotypic difference compared to the WT under the chosen lab conditions, suggesting that their function could be specific to exclusive environmental niches (Fagan & Fairweather, 2014).

Here we focus on *Corynebacterium glutamicum,* an aerobic, Gram-positive soil bacterium that is extensively used in biotechnology and industry for the large-scale biosynthesis of amino acids. Its widespread use is attributed to several advantageous characteristics: biosafety, fast growth to high cell density, genetic stability, absence of autolysis under growth-arrested conditions, low protease activity and a broad spectrum of carbon source utilization (Lee, Na, Kim, Lee, & Kim, 2016). In addition to its growing industrial interest, *C. glutamicum* has emerged as a model organism of the order Corynebacteriles, a subgroup of Actinobacteria that includes important human pathogens, such as *Mycobacterium tuberculosis* and *Corynebacterium diphtheriae*. This taxonomic group exhibits specific characteristics distinct from other bacterial model organisms. Two actinobacterial features of interest are the presence of a multi-layered cell wall that includes an outer membrane mainly composed of long hydrocarbon chains of mycolic acids either esterified to trehalose or attached to the arabinogalactan polymer, which is, in turn, linked to the peptidoglycan meshwork (Fig 1.a) (Dulberger, Rubin, & Boutte, 2019); and a unique polar growth mode, characterized by the insertion of new peptidoglycan at the poles, driven by the coiled-coil elongasome scaffold DivIVA (Letek et al., 2008). This contrasts with model bacilliform organisms such as *Escherichia coli* and *Bacillus subtilis*, which grow laterally by inserting peptidoglycan along the cell wall guided by the actin homologue MreB (Garner et al., 2011)(Fig 1.b).

**Figure 1.**
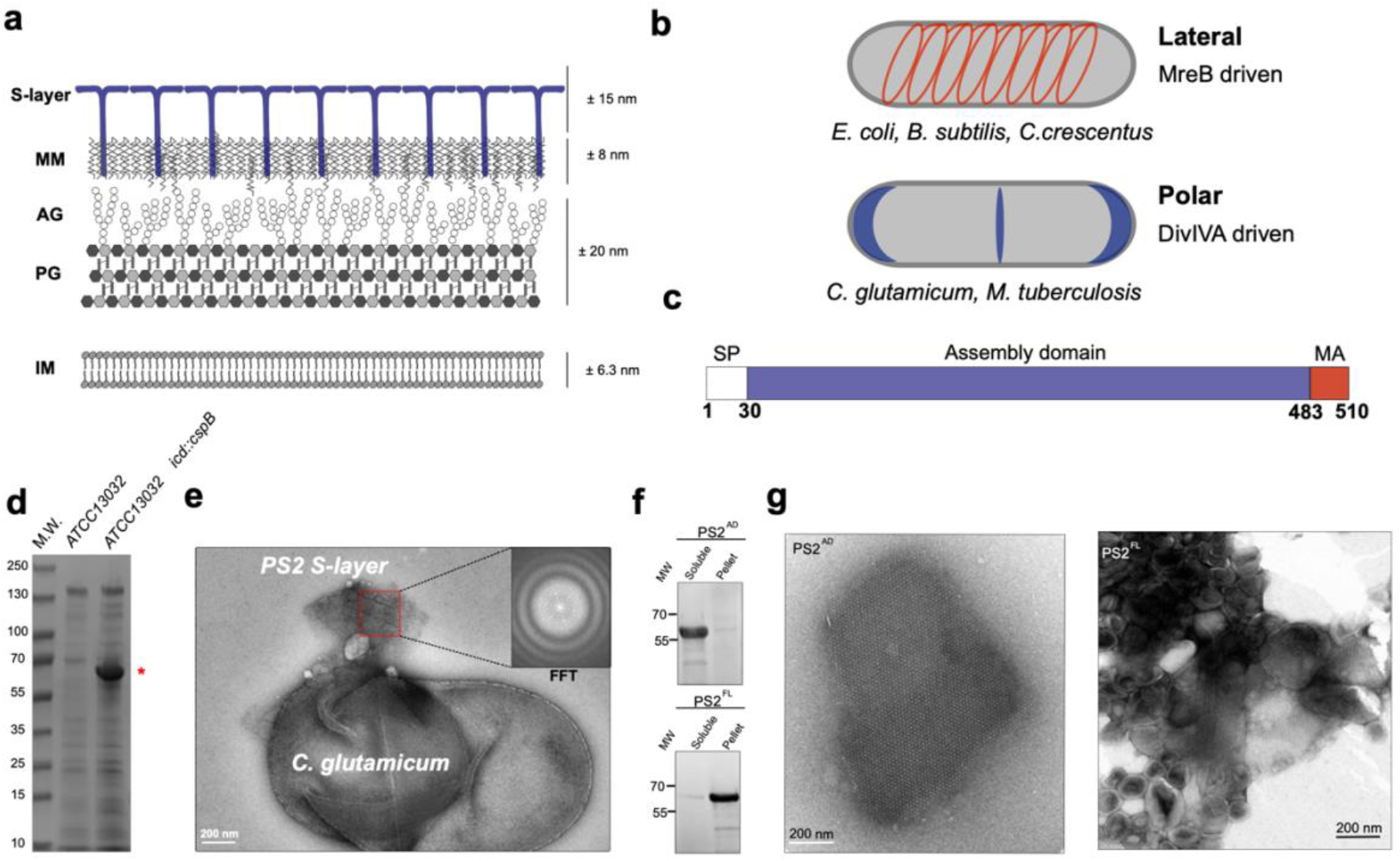
S-layer characterisation and domain organisation of PS2 S-layer from *C. glutamicum*. **a**. Schematic representation of the cell surface organisation of *C. glutamicum*. IM (Inner membrane); PG (peptidoglycan); AG (arabinogalactan); MM (Mycomembrane). **b**. Comparison of the growth modes of Actinobacteria (polar) and model organisms *E. coli, C. crescentus* and *B. subtilis* (lateral). Blue and red represent the localization of DivIVA and MreB respectively which in turn determine PG synthesis. **c**. Domain organisation of PS2. SP (signal peptide); MA (membrane anchor). **d**. SDS- PAGE of the SDS extracted cell surface protein. The strain expressing PS2 under its native promoter expresses large amounts of PS2 (indicated with *). **e**. Negative- stained *C.glutamicum* cells expressing PS2 washed with 0.05% SDS. Detached patches of the PS2 S-layers were observed. (Inset) Fast Fourier Transform of the *ex- vivo* PS2 S-layer. Estimated unit cell parameters are α = β = 171.8 Å and γ = 60°. **f**. Western blot analysis (using anti-HisTag antibody) of recombinant PS2 fractionation; assembly domain (AD) and full-length (FL). **g**. Negative-stained TEM micrograph of *in vitro* reconstituted recombinant soluble PS2^AD^ (left) and lipid-associated PS2^FL^ (right). The scale bar is 200 nm.

On top of the mycomembrane, some Corynebacteriales species display an S-layer that coats their entire cell surface (Fig 1.a). The most studied S-layer protein of this group is the PS2 from *Corynebacterium glutamicum* (Peyret et al., 1993),(Bayan, Houssin, Chami, & Leblon, 2003) whose network has P6 symmetry (Scheuring et al., 2002). PS2 is the product of the *cspB* gene (Peyret et al., 1993) and represents the major secreted protein of the cell (Joliff et al., 1992),(Hansmeier et al., 2004). Early research revealed that PS2 exhibits an abundance of hydrophobic amino acids mainly located in the two terminal regions, identified as an N-terminal signal peptide and a C-terminal cell wall anchoring domain (Chami et al., 1997) (Fig 1.c). As the PS2 S-layer can be stripped from the cell surface by using various detergents it is thought that hydrophobic interactions play a major role in cell wall anchoring. A mutant lacking the 27 C-terminal residues was unable to form an organised S-layer and PS2 was mainly released into the medium (Chami et al., 1997). The current model thus suggests that the C-terminus of PS2 serves as the membrane anchor (MA), tethering the protein to the mycomembrane, which in turn acts as a matrix for 2D crystallization (Bayan et al., 2003) (Fig 1.c). This anchoring mechanism appears to be unique to *Corynebacteriales*, as other Gram-positive bacteria exploit different strategies for S-layer attachment to the cell wall where the S-layer binds directly to the peptidoglycan layer through specialized domains such as the S-layer homology domain (SLH) or the cell wall binding domains (CWB) (Willing et al., 2015),(Blackler et al., 2018). In well-studied bacterial S-layers, the crystallinity of the S-layer is essential to the function (Fioravanti et al., 2019), (Fioravanti et al., 2022),(Sogues et al., 2023) and growth of the S-layer lattice occurs in spatiotemporal coordination with cell elongation and cell wall synthesis (Herdman et al., 2024), (Oatley, Kirk, Ma, Jones, & Fagan, 2020), (Comerci et al., 2019). Where known, S-layer expansion in bacilliform bacteria occurs by the addition of newly exported subunits onto the edge of the lattice, localized at mid-plane. How S-layer expansion and attachment are adapted to the unique cell envelope features of Corynebacteriales is largely unknown.

Here, we present an in-depth analysis of the atomic structure and organization of the PS2 S-layer of *C. glutamicum*. Guided by the lattice structure, we engineered a PS2 variant capable of covalently binding proteins of interest both *in vivo* and *in vitro*, making this S-layer a viable target for its use in biomaterials. Using this engineered PS2 and pulse-chase fluorescent labelling, we tracked its assembly *in vivo*, revealing that this process occurs exclusively at the cell poles. Additionally, an extensive phylogenetic analysis uncovered its scattered distribution within *Corynebacteriales*, along with various genomic contexts, suggesting its paraphyletic distribution and dispersive genome context result from multiple recombination and gene deletion events.

## Results

### Native isolation and purification of recombinant PS2 S-layer

*C. glutamicum* ATCC13032 is widely used in biotechnology and commonly regarded as a reference strain. The strain lacks a 5.97 kb region that contains the *cspB* gene coding for the PS2 S-layer, along with six additional ORFs unrelated to the S-layer biogenesis (Hansmeier et al., 2006). Analysis of this region revealed the presence of a 7 bp direct repeat that could have led to a recombination event responsible for the loss of these genes compared to S-layer containing strains like ATCC4067. In our work, we used the reference strain *C. glutamicum* ATCC13032 in which the *cspB* gene (*Cgl2005*) (Peyret et al., 1993) was inserted under its native promoter into the chromosomal *icd* (isocitrate dehydrogenase) locus, hereafter referred to as ATCC13032 *icd::cspB*. Using SDS-PAGE analysis of surface-extracted proteins, we observed a highly expressed band absent in the wild-type ATCC13032 strain, which corresponded to the expected size of PS2 (Fig 1.d). Negative-stain electron microscopy (ns-EM) inspection of cleared SDS extractions (Chami et al., 1997) of ATCC13032 *icd::cspB* showed the presence of S-layer-like fragments and sheets (Fig 1.e). The power spectrum of the isolated sheets revealed unit cell dimensions of α = β = 171.8 Å and γ = 60°, consistent with previously reported lattice dimensions of PS2 S-layers (Scheuring et al., 2002),(Johnston, Isbilir, Alva, Bharat, & Doye, 2024), thus confirming ATCC13032 *icd::cspB* cells expressed and assembled SDS resistant S-layers. An earlier study showed that PS2 devoid of its membrane anchoring C-terminal domain released monomers into the medium and failed to form an organized S-layer, although proteolytic removal of the C-terminal domain from pre-assembled WT PS2 S-layers did not disrupt the 2D crystal organization (Chami et al., 1997). To further explore whether the C-terminal domain is necessary for S-layer assembly, we cloned the assembly domain (AD) of PS2 (PS2^AD^; residues 30 to 483) in which we replaced the membrane anchoring domain by a hexahistidine tag and overexpressed in the cytoplasm of *E. coli*. Following cell lysis and centrifugation, we observed a pelleted fraction with a gel-like consistency, composed of PS2 as assessed by anti-His western blotting (Fig 1.f). ns-EM confirmed the presence of PS2 S-layer fragments with identical lattice parameters as the *ex-vivo* S-layer (Fig 1. g). We next expressed the full-length variant (PS2^FL^) in the *E. coli* cytoplasm (Fig 1.f), resulting in an insoluble fraction with PS2 lattice characteristics, but associated with vesicle-like structures (Fig 1. g). This suggests that the presence of the MA domain results in a lipid-binding characteristic, consistent with previous observations where the PS2^MA^ domain is responsible for the interactions with the mycomembrane (Bayan et al., 2003).

### *Ex-vivo* cryo-EM structure of the PS2 S-layer from *C. glutamicum*

To obtain high-resolution structural information on the PS2 S-layer, we purified *ex-vivo* S-layer fragments from *C. glutamicum* ATCC13032 *icd::cspB* (see methods). Cryo-EM micrographs of the isolated S-layer predominantly displayed top views of single 2D sheets with planar hexagonal symmetry (Fig 2.a). Determining S-layer structures by cryo-EM represents a challenge as side views are scarce or sometimes non-existent depending on the size of the S-layer fragments. Given the absence of clear side views, we collected tilted images at angles of 15° and 30° allowing for a 3D reconstruction by leveraging the C6 symmetry. The resulting 2D class averages revealed a central hexameric core with six arms, each one extending to two other hexamers, forming a trimeric interface (Fig 2.b). Using single-particle cryo-EM workflow with C6 symmetry, the resulting map reported an average resolution ranging from 2.5 Å to 3.8 Å for various orientations. This difference in resolution is attributed to the uneven representation of views, with the lowest resolution orthogonal to the S-layer plane, corresponding to the underrepresented side views (Supplementary Fig. 1). Despite residual missing wedge artefacts along the Z-axis, the reconstructed map allowed unambiguous docking of the AF2 model of PS2 and further manual refinement in real-space to complete the atomic model (Supplementary Fig. 2). The map reveals a view of the S-layer arrangement and 6-fold symmetry with the presence of six helices forming a conical coiled-coil bundle that extends downward (Fig 2.c,d). The atomic model of PS2 shows that the assembly plane is composed of two distinct interfaces. A C6 hexameric interface where six promoters interact to form the central hexamer, and a C3 trimeric interface formed by the arms of three distinct hexamers in an arm-over-arm arrangement (Fig 2.e). This configuration aligns with the previous Saxton and Baumeister classification as an M6C3 S-layer (Saxton & Baumeister, 1986),(Scheuring et al., 2002). The side view reveals that from each hexamer, six long α-helices extend downwards forming a coiled-coil bundle with a conical shape (Fig 2.e). A single PS2 promoter is composed of eight α-helices adopting an overall banana shape that can be divided into a core and an arm depending on their involvement in the hexameric or trimeric interface respectively (Fig 2.f). The core consists of a 28- residue-long N-terminal random coil and six α-helices tightly packed against each other (H1, H2, H3, H6, H7, H8). Helix H8 extends beyond the core and kinks downwards with an angle of ∼ 120°, where together with the other five protomers they form a hexameric helical funnel. In our structure, we observe clear density until residue A463, leaving the 47 C-terminal residues, which include the predicted membrane anchoring domain, unresolved in the cryo-EM map. The arm region consists of H3, H4, and H5 helices tightly packed by hydrophobic interactions, and connected by two long linkers: one spanning 27 residues and linking H3 to H4, and another of 16 residues forming a distal loop connecting H4 and H5. Notably, H3 is the only helix that contributes to both the core and the arm regions, featuring an insertion by a 10-residue loop. The PS2 S-layer is stabilized by two extensive protein-protein interfaces that make up the C6 contact of the PS2 core and H8 helix, and C3 contacts of the arms (Fig 2.e). The intra-hexameric interface encompasses a total interaction area of 2371 Å^2^, involving the C-terminal H8 helix forming a coiled-coil with hydrophobic knobs-in-holes interactions encompassing 631 Å^2^ (Fig 2.g); and the N-terminal tip of the core 6-helix bundle (i.e. loops H2-H3 and H6-H7) that docks into a large cleft formed in the side of the core region (i.e. H8 and N-terminal coil; (1737Å^2^)) of the neighbouring protomer, at a 60° angle (Fig 2.h). This interaction predominantly involves hydrophilic contacts, including 26 hydrogen bonds and 6 salt bridges (Fig. 2i.). The inter- hexameric contacts are driven by the arm regions and form the C3 trimeric interface. This interface involves the distal loop of one promoter that docks into the cleft formed at the limit between the core and arm regions. This contact presents a total surface area of 1097 Å^2^ involving both hydrophobic and polar interactions (Fig 2.j). Notably, the distal F237 docks into a hydrophobic pocket interacting with F150 from the other protomer, this type of interaction appears to be conserved across several PS2 variants, as many species contain phenylalanine or tryptophan at this position (Supplementary Figure 3). Taken together, the PS2 S-layer is stabilized by a vast network of non-covalent lateral interactions, including hydrophobic contacts, hydrogen bonds, and salt bridges, contributing to its remarkable stability.

**Figure 2.**
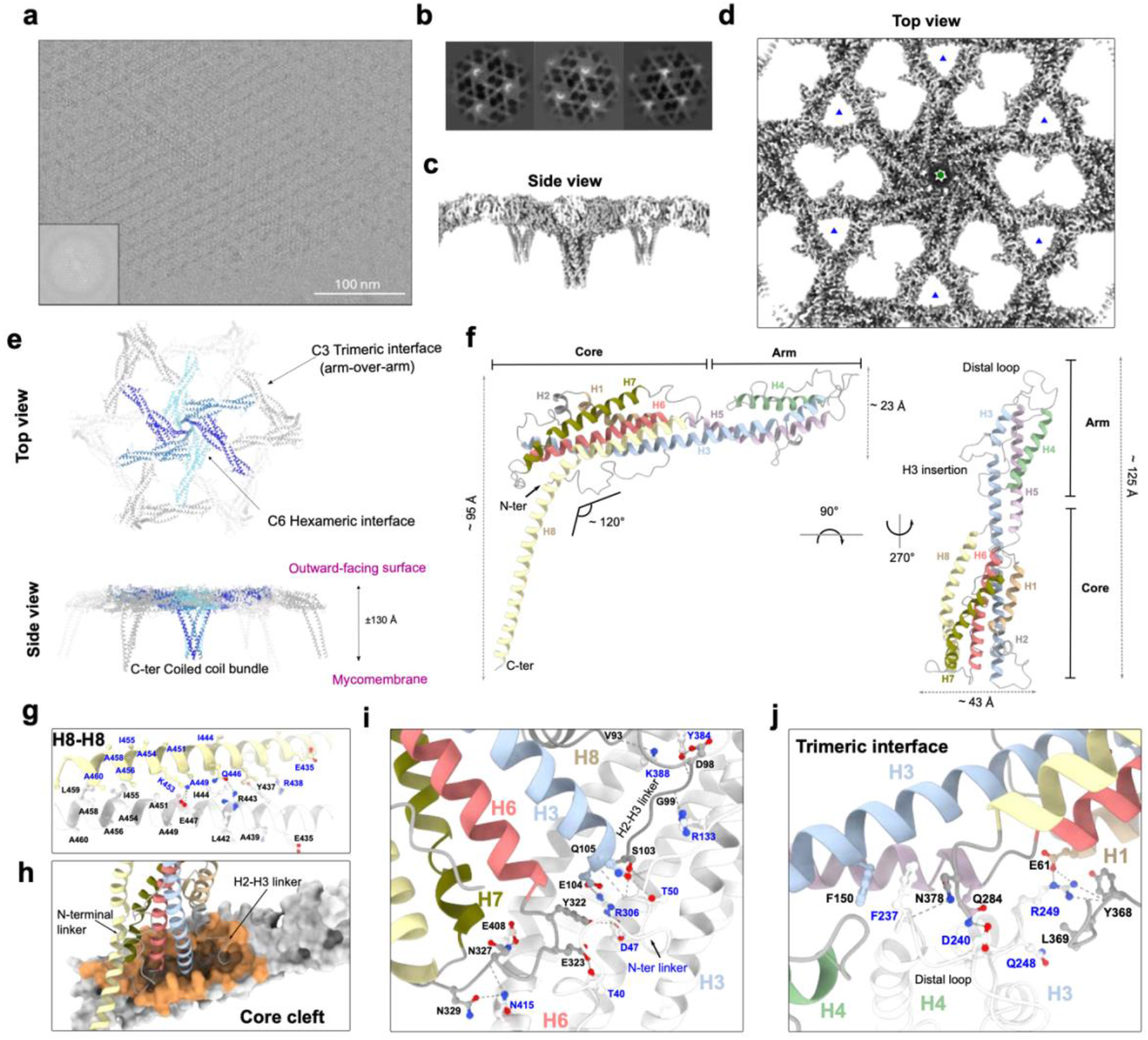
Cryo-EM structure of the *ex-vivo* PS2 S-layer of *C. glutamicum*. **a**. Raw representative Cryo-EM image of a monolayer PS2 S-layer fragment. Scale bar is 100 nm. (Inset) Fast Fourier Transform **b**. Example of 2D class averages of PS2 lattice used for cryo-EM reconstruction. **c, d**. Electron density map after EMReady treatment of the PS2 S-layer shown from the side and top with six-fold symmetry. The average resolution of the map is estimated at 2.51Å. Hexagonal symmetry (green hexagon) and trimeric symmetry (blue triangle) axes are marked. **e**. The atomic model of the PS2 S-layer is shown in ribbon representation, showing both top and side views. The central hexamer is coloured in blue, while the subunits of the interacting hexamers are depicted in grey. Trimer C3 and hexameric H6 interfaces are indicated. **f**. The atomic model of the PS2 monomer, as found in the PS2 lattice, is shown in ribbon representation. PS2 is an all-helix structure, with each helix individually coloured. The PS2 monomer can be divided into two regions: the "core" and the "arm." The "core" forms part of the hexameric interface, while the "arm" forms part of the trimeric interface. The distal loop and H3 insertion loop are shown. **g, h, i, j**. The S-layer lattice is stabilized by three protein-protein interfaces. One monomer is coloured as in panel (**f**) with black labelled residues, while the interacting monomer is shown in white with blue labelled residues. The C-terminal region, formed by H8, creates a 6-helix coiled- coil bundle that stabilizes the hexamer primarily through hydrophobic interactions (**h**). The hexameric interface is stabilized by buried polar interactions formed by the proximal loops that dock into a cleft found in the core region (**h**), involving a vast network of hydrogen bonds representing the largest interaction interface (**i**). Finally, The trimeric interface involves the distal loop, which docks in a cleft at the junction between the arm and core regions (**j**).

### PS2 S-layer properties and functional aspects

Analysis of its surface electrostatics reveals that PS2 is a highly negatively charged protein with 18.7% of its residues being Asp and Glu (pI = 4.21) and located at the protein’s surface (Fig 3.a). We noticed that the outward-facing surface of the S-layer is more negative than the mycomembrane-facing surface, an observation that seems to be recurrent in bacterial S-layers (Baranova et al., 2012),(Lanzoni-Mangutchi et al., 2022). Many bacterial S-layer proteins require structural metal ions for assembly, usually calcium (Baranova et al., 2012), (Sogues et al., 2023), (Herdman et al., 2021). Our cryo-EM map does not suggest the presence of metal ion binding sites in PS2, although the resolution does not allow a fully unambiguous assessment. To assess whether calcium or other divalent ions are required for PS2 stability, we incubated ex-vivo and recombinant PS2^AD^ sheets with 10 mM EDTA. ns-EM analysis showed the presence of S-layers, indicating that divalent metal ions are not required for PS2 stability in pre-assembled S-layers (Supplementary Fig. 4a). Additionally, after unfolding recombinant PS2 S-layers with 8 M urea, isolation of monomers, and refolding in the presence of 10 mM EDTA, ns-EM revealed the presence of S-layer fragments (Supplementary Fig. 4b), showing that divalent ions are not essential for either S-layer assembly or stability.

**Figure 3.**
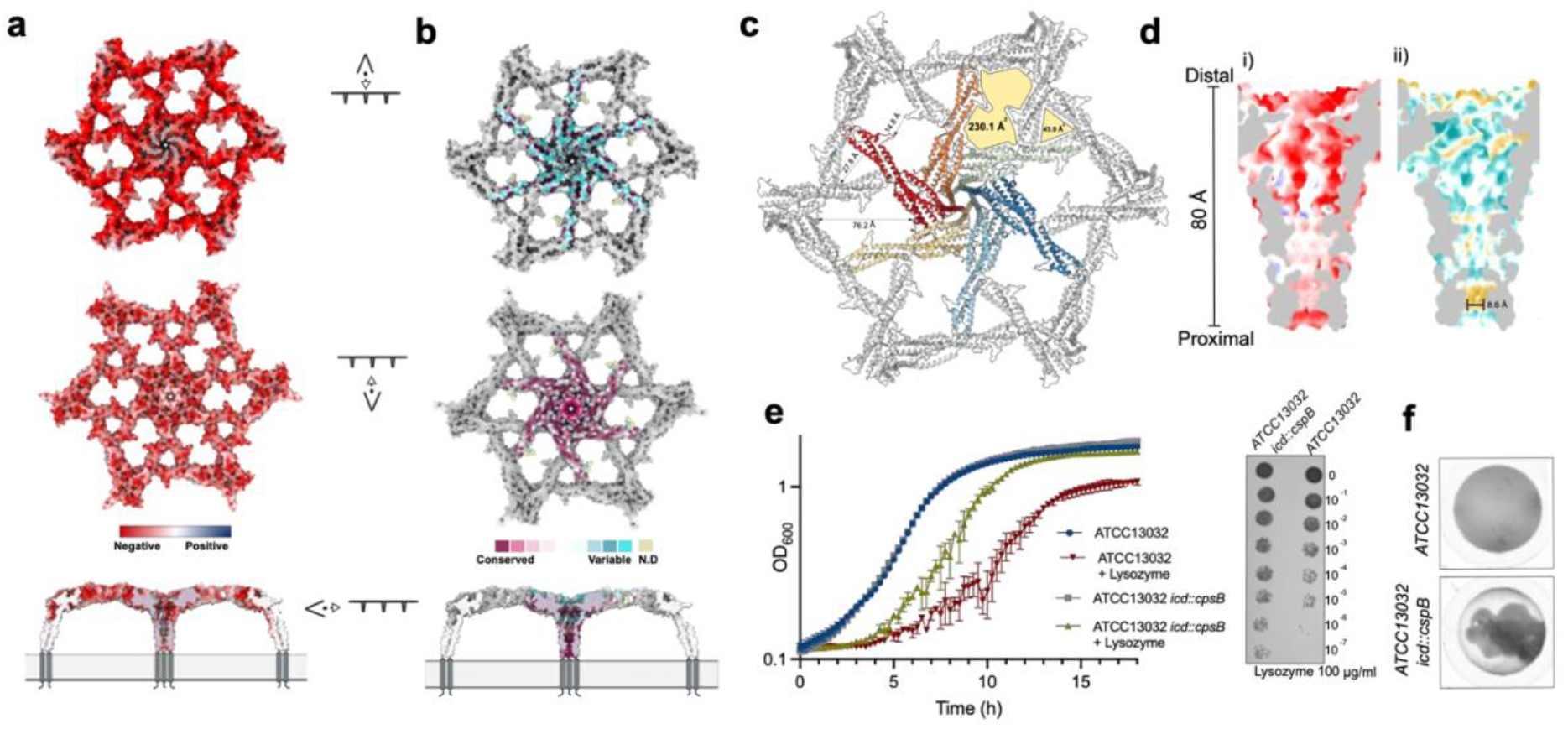
PS2 S-layer is negatively charged and forms large pores. **a**. PS2 hexamer coloured according to its charge distribution (positive in blue to negative in red), shown from extracellular (top), intracellular (middle) and side (bottom) views. **b**. Conservation of the PS2 S-layer mapped on the hexamer (low conservation in blue to high conservation in purple). The extracellular surface (top) is mainly variable, while the intracellular-facing surface (middle) and the inner channel-like (bottom) show higher levels of conservation. **c**. Overall arrangement of the pores in the PS2 S-layer seen from the top. Two main pores are highlighted with yellow filling formed upon S- layer assembly as formed by the C3 trimeric interface and C6 hexameric interface. Areas and distances are indicated in Å. **d**. Cross-section of the channel-like structure formed by the H8 coiled-coil, with the lumen coloured according to charge distribution (i) (as in panel a) and hydrophobicity (ii) from hydrophobic regions in yellow to hydrophilic regions in blue. The narrowest constriction of the pore is indicated **e**. Growth curves (left) comparing ATCC13032 *icd::cpsB* (expressing PS2 (grey and yellow)) with ATCC13032 (red and blue) in the presence (yellow and red) or absence (grey and blue) of 100 μg/ml lysozyme. Growth curve data are sample mean ± s.d., representative of n = 3 biological experiments. Lysozyme was added at the start of the measurements. Lysozyme sensitivity assay (right panel) after incubating ATCC13032 and ATCC13032 *icd::cpsB* in 100 μg/ml of lysozyme overnight. S *icd::cpsB icd::cpsB* amples were normalised to an OD600 of 0.5, serially diluted 10-fold, and spotted onto an LB agar plate. **f**. Different sedimentation properties are observed when ATCC13032 expresses the PS2 S-layer compared to its absence. This is manifested from imaging a single well in a 96-well plate after overnight growth, followed by 12 hours of incubation at room temperature without shaking.

S-layers often show low levels of sequence conservation even within the same phylogenetic group. We analysed the sequence conservation of PS2 and mapped it onto the structure. Strikingly, the external and mycomembrane facing surfaces of the S-layer show increased sequence variation or increased conservation, respectively (Fig 3.b). The positive selection for variation on the external face aligns with the concept that S-layers might evolve rapidly to adapt to new ecological niches or defend against emerging environmental threats. The most conserved part of the protein maps to the C6 hexameric interface and involves the proximal region of the core (H2-H3 linker) and the docking cleft (Supplementary Fig. 5). Similarly, residues that contribute to the trimeric interface are conserved suggesting that the overall architecture and assembly mechanism is the same across different PS2 S-layers. The extended arrangement of PS2 results in two major pores in the lattice (Fig 3.c). The trimeric pore has a diameter of 27.8 Å and an area of 43.9 Å^2^. The largest pore is formed by the interaction of two different hexamers with a maximum diameter of 76.2 Å with a total area of 230.1 Å^2^. This pore shows a constriction of 14.8 Å due to the presence of the H3 insertion loop (Fig 3.c). Finally, a small funnel-like pore is formed by the H8 coiled-coil at the centre of the hexamer, possibly extending into a channel through the mycomembrane. This channel-like pore with a height of ∼80 Å shows a lumen that is mainly negatively charged but becomes more neutral proximal to the cell, where a belt of leucines (L459) and isoleucines (I455) forms a hydrophobic constriction of 8.6 Å in diameter, the narrowest point in the channel (Fig 3.d). As such, the PS2 S-layer can be viewed to represent a continuous semipermeable layer anchored into the mycomembrane on ∼8 nm high pedestals, with plausible roles as selectivity and/or barrier and/or support structure (Fig 3.a, b). To test these hypotheses, we performed a lysozyme susceptibility test, a standard method for assessing cell wall integrity. Growth curves of ATCC13032 and ATCC13032 *icd::cspB* showed no difference in the absence of lysozyme, suggesting that S-layer expression does not affect fitness under laboratory conditions. However, upon lysozyme addition, the culture of the strain lacking the S-layer (ATCC13032) showed reduced growth (Fig 3.e). This result aligns with previous findings (Matsuda et al., 2014) and mirrors a similar result observed in *C. difficile* (Kirk et al., 2017),(Lanzoni-Mangutchi et al., 2022). Given that lysozyme is a small protein with a hydrodynamic radius of 1.9 nm and therefore expected to be capable of passing through the PS2 pores, we hypothesize that the partial protection from the S-layer may result from its adsorbing lysozyme molecules and/or acting as a mechanical support that helps the cell envelope maintain turgor pressure, as seen for *B. anthracis* S-layers (Fioravanti et al., 2019), (Fioravanti et al., 2022), (Sogues et al., 2023). Finally, another phenotypic difference observed while growing ATCC13032 with and without PS2 expression was the variation in coagulation and sedimentation properties. Unlike its wildtype parrent, the ATCC13032 *icd::cspB* strain expressing PS2 exhibited strong flocculation under static growth conditions (Fig 3. f). Thus the presence of the PS2 S-layer appears to alter the surface properties of *C. glutamicum*, resulting in an increased coagulation of cells, a property that may also impact biofilm formation and adhesion to surfaces (D. Zhang et al., 2022).

### Engineering and Biogenesis of the S-layer in *C. glutamicum*

S-layers are attractive biomaterials with interesting properties such as regular auto-assembly. We have engineered the PS2 S-layer of *C. glutamicum* to display the SpyTag, which forms a covalent bond with SpyCatcher-tagged proteins (Zakeri et al., 2012). Structural analysis revealed that both termini of PS2 are not surface exposed suggesting that the SpyTag should be added internally. Guided by the structure of the S-layer lattice, we introduced the 18 residues that form the SpyTag in the H3 insertion loop. This loop is absent in most of the PS2 sequences except for *C. glutamicum* strains ATCC13870 and ATCC14068. In addition, sequence analysis showed that the H3 isertion loop is composed of a sequence repeat at the DNA level resulting in the amino acid sequence: <u>SINPD</u>G<u>SINPD</u>, suggesting that it arose from a small duplication event. Therefore, we inserted the SpyTag in position 168 (PS2^SpyTag^) (Supplementary Fig. 6). To test that the insertion is functional and does not impact PS2 self-assembly, we purified the recombinant PS2^AD-SpyTag^ and mixed it with 1.5 molar excess of SpyCatcher-mCherry. Fluorescence light microscopy showed the presence of fluorescent recombinant S-layers (Fig 4.a). As a control, we used the WT version (PS2^AD^) which did not exhibit any fluorescent signal (Supplementary Fig. 7a). ns-TEM confirmed that the addition of SpyTag in complex with SpyCatcher-mCherry does not alter the formation of ordered 2D sheets (Supplementary Fig. 7b). These observations indicate that engineered PS2^SpyTag^ is functional and allows for specific binding of a SpyCatcher fusion protein. Next, to engineer PS2 *in vivo*, we cloned the PS2^SpyTag^ under its native promoter into the pTGR5 shuttle plasmid (Ravasi, Peiru, Gramajo, & Menzella, 2012). We expressed this construct into the S-layer lacking strain (ATCC13032) and validated its expression (Fig 4.b). To explore *in vivo* labelling of the PS2 S-layer, we incubated *C. glutamicum* ATCC13032 expressing PS2^WT^ or PS2^SpyTag^ with SpyCatcher-mCherry and visualized the cells using fluorescence microscopy. While no significant mCherry signal was observed in the strain expressing the PS2^WT^, the PS2^SpyTag^ strain presented a bright and uniform fluorescence signal surrounding the cell surface (Fig 4.c). These results demonstrate the potential for engineering PS2 to selectively attach specific proteins to the extracellular surface of *C. glutamicum*.

**Figure 4.**
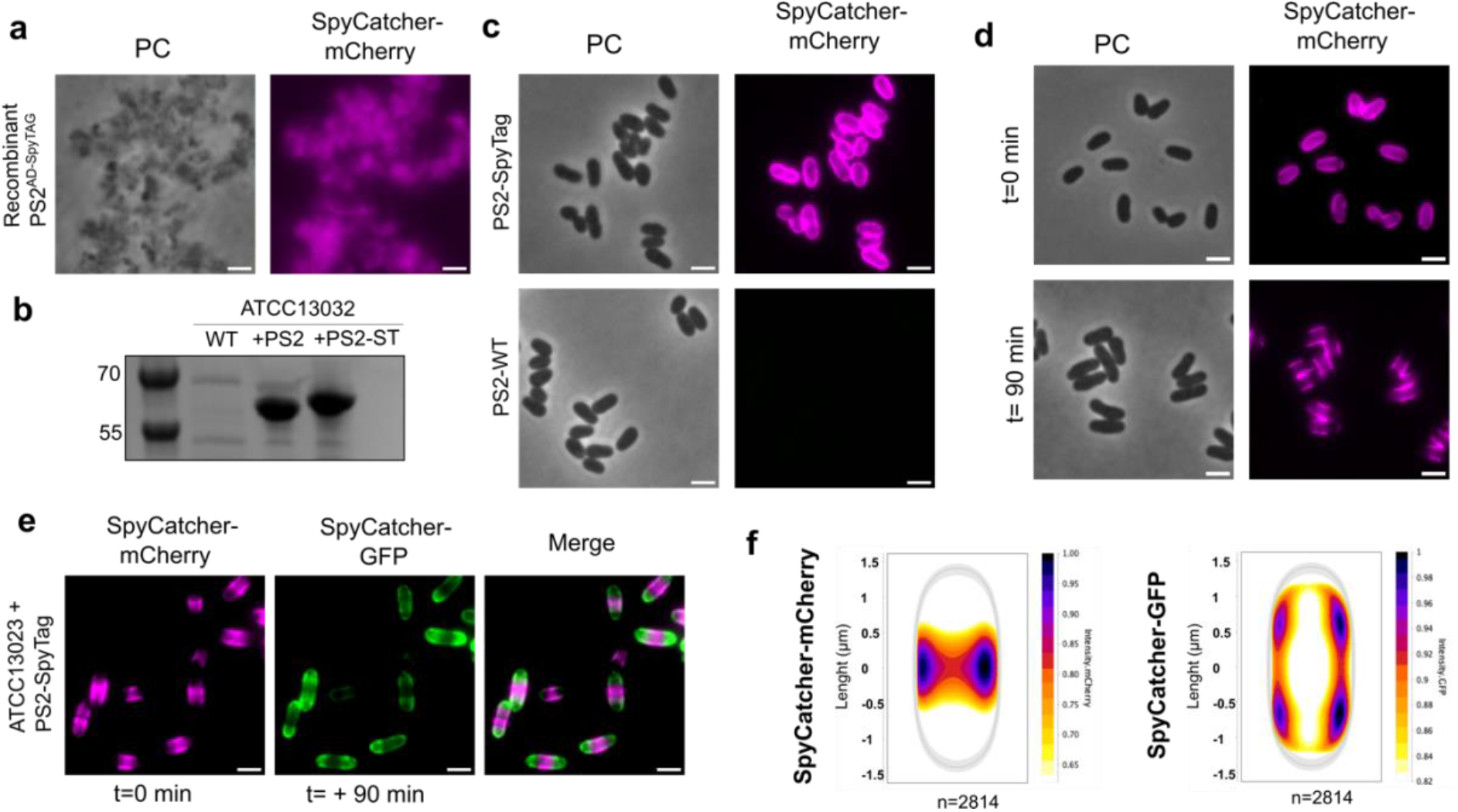
Engineering of the PS2 S-layer and polar assembly. **a**. Micrograph of recombinant PS2^AD-SpyTAG^ incubated with Spycatcher-mCherry shows *in vitro* functional engineered PS2. **b**. SDS-PAGE of the extracted cell surface protein comparing ATCC13032 and transformed strain expressing PS2 and PS2^SpyTAG^. The strain expressing the SpyTag shows an increased molecular weight. **c**. Micrograph of *C. glutamicum* expressing PS2^SpyTAG^ (top) or WT PS2 (bottom) and incubated with SpyCatcher-mCherry. The surface of the strain expressing the engineered PS2 shows a homogeneous fluorescence surface signal indicating the covalent complex formation with SpyCatcher-mCherry. The WT shows no binding. **d**. Micrograph of Spycatcher- mCherry stained PS2^SpyTAG^ at t=0 min and t=+90 min. At a later point, the old (stained) S-layer is restricted to the middle of the cell with no apparent diffusion. **e**. Representative micrograph of PS2^SpyTAG^ expressing strain first strained with Spycatcher-mCherry and pulse chased with a second stain using Spycatcher-GFP after 90 min. The new S-layer (green) is inserted at the poles. In all of the above panels, the scale bar = 2 µm. **f**. Normalized heat map representing the localization pattern of SpyCatcher-mCherry (old S-layer) and SpyCatcher-GFP (new S-layer). A total of 2814 cells were analysed. The images are representative of experiments made independently in triplicate.

Next, we used the ability to covalently label *in vivo* PS2 S-layers to investigate the molecular mechanisms of S-layer biogenesis by means of a two-colour pulse-chase experiment with two distinct fluorescent SpyCatcher fusions. Prior studies in Gram-positive (Oatley et al., 2020), Gram-negative bacteria (Comerci et al., 2019) and Archaea (Farid Abdul-Halim et al., 2020), showed that *de novo* S-layer assembly is localized predominantly at mid-plane, suggesting a co-localization with sites of cell elongation and cell wall synthesis. Interestingly, *Corynebacteriales* grow from their poles, where the cytoskeletal protein DivIVA (also known as Wag31) guides the elongasome and peptidoglycan insertion. The second site of cell wall biosynthesis is the septum where the FtsZ-guided divisome incorporates the new cell wall (Meyer & Bramkamp, 2024),(Sogues et al., 2020) (Supplementary Figure 8). If S-layer assembly co-localizes with peptidoglycan synthesis, we hypothesized that in *C. glutamicum*, S- layer biogenesis would occur primarily at the poles and/or at mid-cell. To test that hypothesis we first, we saturated the surface of *C. glutamicum* ATCC13032 PS2^SpyTag^ with a pulse labelling of SpyCatcher-mCherry and observed that after 90 minutes of continued growth, the poles were devoid of fluorescent signal (Fig 4.d). This result shows that the PS2 S-layer exhibits non-diffusive behaviour, as the old, labelled PS2 is not redistributed across the cell surface. Second, this data suggested that new S- layer is incorporated at the poles. To further support this idea, we performed chase labelling of the new, unstained S-layer with SpyCatcher-sfGFP. The images showed that the old S-layer (labelled with SpyCatcher-mCherry) localized over the lateral body, away from the poles, whereas the newly synthesized S-layer (labelled with SpyCatcher-GFP) primarily displayed a polar signal (Fig 4.e). As expected from the visual inspection of the data, fluorescence signal quantification of >2000 dually labelled cells confirmed these patterns (Fig 4.f). Thus, our results support the hypothesis that new S-layer assembly co-localizes with the zones of polar peptidoglycan synthesis and cell elongation, and suggest that this may be a generic characteristic across different S-layers and different cell elongation strategies.

### Phylogenetic analysis of PS2

The available structures of bacterial S-layers show a lack of structural homology across genera (Fioravanti et al., 2019)(Baranova et al., 2012),(Lanzoni-Mangutchi et al., 2022),(Bharat et al., 2017),(von Kügelgen et al., 2023) indicating that S-layers have arisen independently multiple times throughout evolution (Johnston et al., 2024). Here we set out to study the phylogeny and distribution of PS2 in the order *Corynebacteriales*. Our analysis revealed that PS2 homologues are exclusively found in *Corynebacterium*, suggesting genus specificity. However, its presence is sporadic, with only 102 hits (i.e. cutoff of e-value 1e-05 to ATCC13032 PS2 –GenBank sequence AAX43986.1) out of the 2325 genomes analysed (4.25%) (Fig 5.a, Supporting Data). PS2 genes show a scattered, paraphyletic distribution across *Corynebacterium* species, sometimes specific to just some strains within the same species. This is the case for *C. glutamicum*, where the reference strains ATCC4067 and ATCC13032 respectively hold or lack the *cspB* gene. The presence of a 7 bp sequence repeat, an integrase and an IS element in the genomic region encoding *cspB* suggests that a recombination event may have resulted in the acquisition or deletion of *cspB* in these strains (Hansmeier et al., 2006). Moreover, the paraphyletic distribution of *cspB* could be the result of repetitive losses of *cspB* or acquisitions through horizontal gene transfer. To discriminate between these two scenarios, we studied the genomic context of the *cspB* gene. The result of this analysis showed that *cspB* presents varying genomic contexts across different species, whereas it seems to be conserved within related species (Fig 5.b). Next, we compared the genomic loci where *cspB* is present with those of closely related species lacking PS2 (Supplementary Fig. 9). Our analysis revealed many different scenarios including a substantial number of deletions, insertions, and even inversions surrounding the *cspB* gene. These genomic alterations sometimes affected only the *cspB* gene itself or involved neighbouring genes, suggesting that *cspB* gene resides within or is linked to mobile genetic regions. Nevertheless, the phylogenetic tree of the *cspB* gene appears to largely follow the full genome phylogeny of the *cspB* positive strains (Supplementary Fig. 10), suggesting that the paraphyletic distribution and dispersed genomic context of PS2 are likely a result of multiple recombination and gene deletion events.

**Figure 5.**
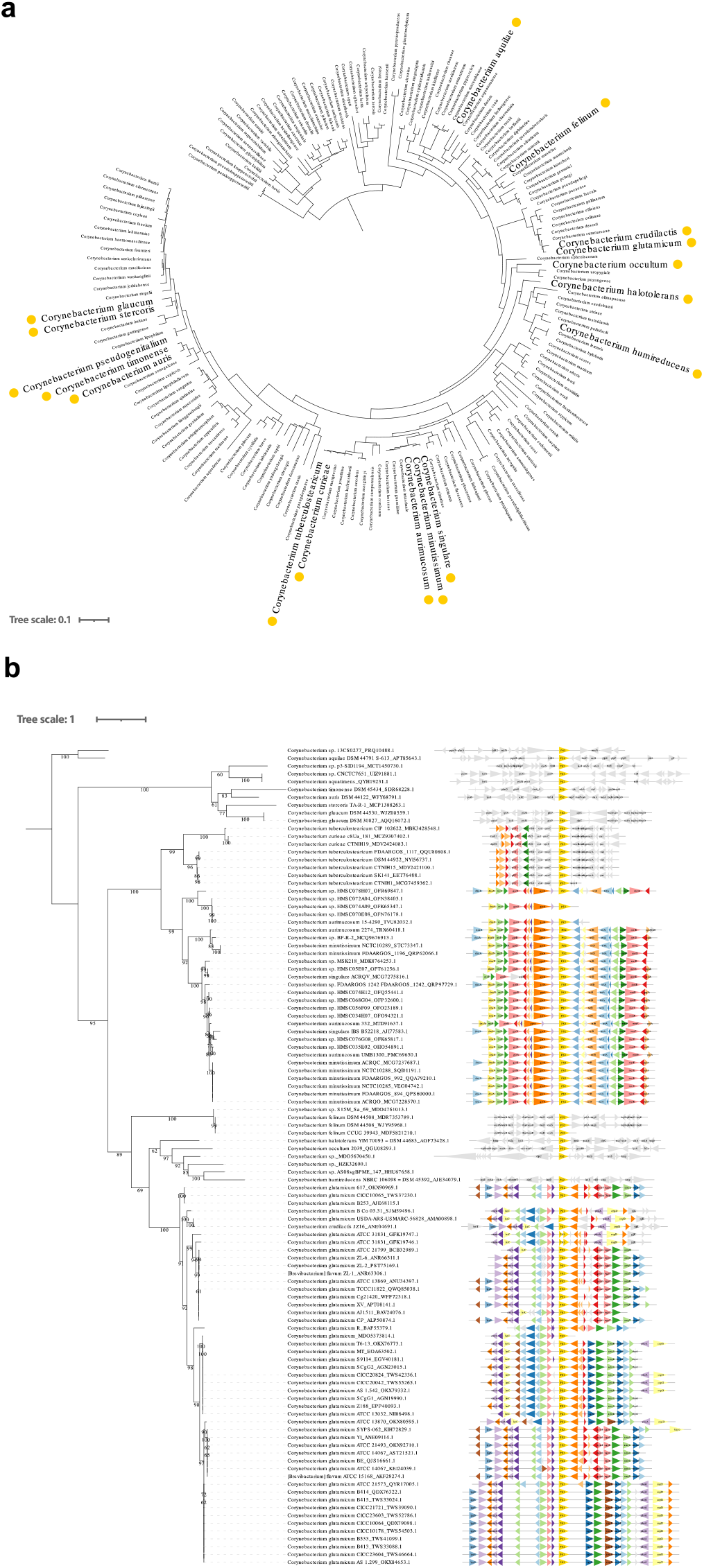
PS2 phylogeny and genome context show multiple recombination events. **a**. Phyletic pattern of the presence of PS2 in a reference phylogeny of the *Corynebacterium* genus. Yellow circles indicate that at least one strain for that species codes for protein PS2. **b**. Phylogenetic tree of protein PS2 (centred in yellow) and genomic context. Triangles in colours correspond to genes frequently found in the same locus as *cspB* (PS2). Triangles without labels correspond to genes of unknown function. Numbers on branches correspond to ultrafast bootstrap supports.

## Discussion

*Corynebacterium glutamicum* has emerged not only as a biofactory but also as a model organism in Actinobacteria, one of the largest bacterial phyla on Earth. This study provides new structural and biological insights into one of the most complex bacterial cell envelopes. Atop the cytoplasmic membrane, the *Corynebacterium* cell envelope has a peptidoglycan (PG) layer that is decorated with arabinogalactan (AG) oligosaccharides (Fig. 1a), with a combined height of about 20 nm. The heavily cross- linked PG layer provides the main mechanical support to the cell envelope, requiring a strongly coordinated assembly to assure cell envelope integrity throughout cell growth and division (Meyer & Bramkamp, 2024). Unique to Actinobacteria, a second membrane is found composed of mycolic acids. This mycomembrane (MM) creates an additional, amphiphilic permeability barrier similar in principle, but structurally distinct, to the outer membrane (OM) in diderm bacteria. Together with the cytoplasmic membrane, the MM delineates a periplasmic space of >20 nm. As for diderm bacteria, the presence of the mycomembrane requires dedicated transport pathways for the export and uptake of proteins and metabolites, operated by means of beta-barrel pores and large oligomeric complexes (Viljoen et al., 2017). Finally, in some Corynebacteriales species and strains, the mycomembrane is additionally covered by an S-layer. Here, the 3D cryoEM structure of extracted PS2 S-layers from *C. glutamicum* reveals this S-layer as a continuous, semi-porous monolayer of C6:C3 symmetry and 25 Å thickness, with a regular network of gaps of ∼27 Å and ∼76 Å maximum diameter (∼44 Å^2^ and ∼230 Å^2^ surface area, resp.; Fig 2b, 3c). This monolayer is formed by PS2 hexamers with protruding ‘arms’ that maintain C3 contacts with neighbouring hexamers. The PS2 hexamers are anchored in the mycomembrane by the C-terminal ∼27 residues of the protein (absent in the reported structure), which are found at the end of a funnel-like coiled-coil of ∼70 Å height and formed by the elongated H8 helix. As such, PS2 hexamers attain a parasol-like structure (Fig 2c, e), resulting in the formation of a 7 nm pseudoperiplasmic space atop the mycomembrane. The functional significance of this pseudoperiplasmic space, or indeed the PS2 S-layer remain largely unknown.

In the presence of the S-layer, we find *C. glutamicum* became less sensitive to extracellular lysozyme. However, the porous structure of the PS2 lattice is compatible with the passage of proteins of up to 50-100 kDa (i.e. ∼50 to 60 Å diameter when considered spherical and average density of 1.35 g/cm^3^; (Fischer, Polikarpov, & Craievich, 2004), making it unlikely that PS2 would act as a physical barrier to most lytic enzymes. In addition, most enzymes, like lysozyme, would need to pass at least the mycomembrane to reach their targets. How then does PS2 protect from lysozyme activity? Possibly, PS2 could still lower lysozyme infiltration by means of electrostatic repulsion or absorption onto the S-layer. Alternatively, the S-layer may provide a mechanical support to the mycomembrane and corynebacterial cell envelope that helps protect it from osmotic lysis in case of a weakened PG cell wall. Such mechanosupportive function has been demonstrated at least for the Sap and EA1 S-layers in *Bacillus anthracis* (Fioravanti et al., 2022),(Sogues et al., 2023), and is a main function of archaeal S-layers (Albers & Meyer, 2011).

To maintain its barrier and mechanical function, the secretion and assembly of cell envelope components need to be coordinated with cell growth and division. These cellular processes are orchestrated by cytoskeletal proteins that act as recruitment signals and provide the dynamics of cell cycle progression. The mechanism by which the S-layer assembly expands in coordination with the entire cell envelope remains unclear. This is particularly puzzling given that S-layers form regular lattices, which typically grow by the addition of subunits at their edges. Therefore, unless the S-layer is composed of a mosaic of crystalline microdomains or the cells are capable of dynamically assembling and disassembling the lattice, the S-layer would be expected to associate with the cell envelope as a continuous unit, with lattice edges exposed only in regions of cell expansion. Only recently a handful of studies have shed light on the dynamics of the S-layer during cell growth. In bacteria, this process has been studied in *Caulobacter crescentus* (Comerci et al., 2019) and *Clostridioides difficile* (Oatley et al., 2020), revealing that S-layer growth occurs mainly at mid-cell, indeed coinciding with the regions where new peptidoglycan is inserted by the divisome machinery. In *C. crescentus*, inhibition of MreB (a major cytoskeletal component that drives the elongasome) resulted in delocalised S-layer insertion (Herdman et al., 2024). Corynebacteriales lack MreB and instead contain the coiled-coil elongasome scaffold DivIVA, which is responsible for their rod shape and involved in localising the peptidoglycan synthesis machinery to the poles (Letek et al., 2008). Our study is the first to examine S-layer biogenesis in polar-growing bacteria, revealing that this process occurs exclusively at the poles. We did not observe new S-layer being added at mid-cell where divisome-driven peptidoglycan synthesis also takes place, indicating that S-layer assembly is only associated with the elongasome. Previous models for S- layer assembly suggest that cell wall expansion is a driving force in cell envelope growth and a predictor of local S-layer biogenesis, where a pool of free S-layer proteins (SLPs) exists in the cell wall to plug S-layer-free regions (Herdman et al., 2024), (Oatley et al., 2020), (Comerci et al., 2019). If this was true for *Corynebacterium*, we would also observe labelling of the new S-layer at mid-cell, at least in cells initiating division. Different to most bacterial S-layers, however, the *Corynebacteriales* S-layers are membrane anchored (i.e. the mycomembrane) rather than attached to the cell wall or a secondary cell wall polymer. This physical separation and the fact that we do not observe PS2 S-layer growth at mid-cell suggests the existence of a new *Corynebacteriales* model for the delivery and assembly of S-layer subunits. Likely, the PS2 S-layer acts as a continuous lattice floating on the mycomembrane by means of the C-terminal transmembrane anchors. New subunits are added at the elongating poles, where expansion of the cell wall and mycomembrane result in the exposure of the lattice edge of the S-layer. Interestingly, polar growth leads to a phenomenon known as the divisome-elongasome transition (Martinez et al., 2023). After cell division, the septum transforms into a new pole that requires the assembly of a new elongasome. So, our observations are compatible with a model where S-layer growth at the division plane, only occurs after the divisome-elongasome transition, and may suggest that S-layer secretion and/or biogenesis are directly or indirectly coordinated by elongasome components. The PS2 S-layer is exported by means of an N-terminal leader sequence and the SEC translocon. How and where it traverses the mycomembrane to reach the cell surface is unknown. Possibly, the elongasome scaffold orchestrates these sites of export and secretion. Even so, our demonstration that the recombinant introduction of merely *cspB* into ATCC13032, as well as our phylogemic analysis showing that the horizontal acquisition of just *cspB* is sufficient for functional PS2 assembly, suggest that PS2 secretion does not require dedicated machinery, but occurs through a common, pre-existing pathway. Future studies will be required to identify the mode of secretion, and to evaluate if and how secretion of PS2 is coordinated by the elongasome. Such studies could focus on (i) the localization of the SEC machinery used by PS2 for translocation across the inner membrane (Houssin, Nguyen, Leblon, & Bayan, 2002) (ii) tracking new S-layer formation under DivIVA depletion conditions, (iii) evaluating S-layer assembly in null mutants of known mycomembrane insertion of translocation pathways and (iv) localizing a monomeric PS2 (assembly-incompetent mutant) to determine if it diffuses in the cell envelope or exhibits polar localization.

Finally, S-layers have attracted significant interest in bioengineering materials and synthetic biology as a display platform due to their crystalline self-assembly behaviour which facilitates precise spatial positioning and high-density material display. Our work demonstrates that recombinant introduction of *cspB* into *C. glutamicum*, and likely other species, readily results in the secretion and assembly of PS2 S-layers. Moreover, we find that the PS2 S-layer alters cell coagulation and flocculation, properties important for fermentation and downstream processing behaviour. Furthermore, we show that the engineering of PS2 by insertion of the SpyTag, results in an easy platform for covalent surface display both *in vitro* and *in vivo* by means of SpyCatcher-SpyTag and related protein conjugations technologies. Our structural analysis predicts that likely, these same sites (i.e. the H3 insertion loop, residues 162- 171) are amenable to the insertion and abundant, regular surface display of larger fusion peptides or whole proteins. As such, we anticipate that this structural work and PS2 engineering technology provide an interesting expansion to the broad use of *C. glutamicum* as an industrial workhorse for the production of (poly)peptides, amino acids and other fermented materials.

## Methods

### Bacterial strain and growth conditions

All bacterial strains used in this study are listed in Supplementary Table 1. *Escherichia coli* DH5α was used for cloning purposes and was grown in LB media or agar plates at 37 °C supplemented with 50 µg/ml kanamycin or 50 µg/ml Ampicillin when required. For protein production, *E. coli* BL21 (DE3) was grown in TB media supplemented with 50 µg/ml Ampicillin at the appropriate temperature for protein expression.

*Corynebacterium glutamicum* ATCC13032 was used as a wild-type (WT) strain and *C. glutamicum* ATCC13032 *icd::cpsB* expressing the *cspB* gene (Cgl2005) (Peyret et al., 1993). *C. glutamicum* strains were grown in LB or BHI media at 30 °C and 120 rpm and were supplemented with 25 µg/ml kanamycin and/or 20 ml/L of Sodium lactate when required to induce higher levels of PS2 expression (Soual-Hoebeke et al., 1999).

### Cloning for recombinant production in *E. coli*

The *cspB* gene (Uniprot ID: Q04985) coding from residues 30 to 510 was amplified by PCR using oligos p849 and p868 whereas the assembly domain construct (PS2^AD^) (residues 30 to 483) was amplified with oligos p849 and p850 using as a template the gDNA of the *C. glutamicum* ATCC13032 *icd::cpsB*. The PCR fragments were cloned into a linearised pASK-IBA3plus vector (using primers p321 and p322) by Gibson assembly leading to plasmids A232 and A231 and transformed into chemically competent DH5α *E. coli* (New England BioLabs). PS2-spyTAG versions were produced by site-directed mutagenesis using oligos p873 and p874 leading to plasmid A235. SpyCatcher-GFP and SpyCatcher-mCherry were synthetically ordered (Integrated DNA technologies - IDT) and cloned into the linearised pASK-IBA3plus vector leading to plasmids A233 and A234 respectively. Plasmids were sequence verified (Eurofins). All plasmids and primers are listed in Supplementary Table 1.

### Cloning for recombinant protein expression in *C. glutamicum*

For ectopic expression of PS2 variants in *C. glutamicum* ATCC13032, we amplified the *cspB* gene with its native promoter with oligos p864+p865 using as a template the ATCC13032 *icd::cpsB* (which contains the *cspB* gene with its native promoter). PCR fragment was cloned by Gibson assembly into a linearised pTGR5 (using oligos p862+p863) leading to the formation of the A236 plasmid. Insertion of the SpyTAG was done using oligos p873+p874 and A236 as a template, leading to the A242 plasmid. Plasmids were sequence verified (Eurofins) and transformed into electrocompetent *C. glutamicum* cells as described in (Sogues et al., 2020).

Sequences of interest are found in Supplementary Table 2.

### *Ex-vivo* PS2 purification

To purify ex-vivo PS2 S-layer fragments from C. glutamicum ATCC13032 icd::cpsB, we grew 500 ml of culture in LB medium with 2% sodium lactate overnight at 30°C. The culture was harvested by centrifugation (10 min at 5000 x g), and the pellet was resuspended in PBS + 1% SDS, followed by a 2-hour incubation with shaking. The mixture was then homogenized using a blender and loaded onto a 20% sucrose cushion. After centrifugation (30 min at 4800 x g), the layer above the cushion, enriched with S-layer fragments, was recovered while cells were found in the pellet.

The S-layer fragments were centrifuged again (30 min at 20,000 x g) and washed with 100 mM NaCl and 20 mM Hepes pH 7.

### Growth curves and lysozyme resistance

All strains were initially plated on LB agar for 2 days at 30°C. A single colony from each plate was then inoculated into LB media supplemented with 2% sodium acetate and incubated overnight at 30°C with 120 rpm shaking. The following day, 2 ml of LB media supplemented with 2% sodium acetate were inoculated with the overnight cultures to achieve a starting OD600 of 0.05, and 200 μl of each culture was dispensed into individual wells of a 96-well plate. For experiments involving lysozyme, a final concentration of 100 μg/ml was used. The 96-well plates were then loaded into the Cytation One system (BioTek) and incubated at 30°C with double orbital shaking. OD600 measurements were recorded every 15 minutes. Data analysis and plotting were performed using Prism8 software. All experiments were conducted in triplicate, and the results are presented as the mean ± standard deviation.

### Protein expression and purification

PS2 and SpyCatcher derivatives were expressed in *E. coli* BL21 (DE3) grown in Terrific Broth (TB) supplemented with 100 µg/ml of Ampicillin at 37 °C and induced with 200 µg/L anhydrotetracycline when OD600 reached 0.6. Following induction, the temperature was dropped to 23°C for overnight expression. Next day, cells were harvested by centrifugation (20 min at 5000 xg) and pellets were kept at −20 °C. Cell pellet was resuspended in 100 ml of lysis buffer (50mM HepespH8,300mM NaCl,1mM MgCl2, DNase, lysozyme and EDTA-free protease inhibitor cocktails (ROCHE)) at 4 °C and lysed by sonication. The lysate was centrifuged for 60 min at 30,000 × g at 4 °C. For SpyCatcher derivatives, the cleared lysate was loaded onto a Ni-NTA affinity chromatography column (HisTrap FF crude, GE Healthcare) and washed extensively with buffer A (50 mM Hepes pH8, 300 mM NaCl, 10 mM imidazole). His-tagged proteins were eluted with a linear gradient of buffer B (50 mM Hepes pH8, 300 mM NaCl, 5% glycerol, 1 M imidazole). The eluted fractions containing the protein of interest were pooled, concentrated and loaded onto a Superdex 75 16/60 size exclusion (SEC) column (GE Healthcare) pre-equilibrated at 4°C in SEC Buffer (50 mM Hepes pH8, 150 mM NaCl). The peak corresponding to the protein was concentrated, flash-frozen in small aliquots in liquid nitrogen and stored at -80°C. For PS2^AD^, following centrifugation of the lysate, an additional layer with a gel-like consistency was observed between the supernatant and the pellet, primarily containing PS2. This PS2-containing pellet was carefully collected and subsequently resuspended in SEC buffer supplemented with 1% DDM (n-dodecyl-β-D-maltoside), followed by overnight incubation. The next day, PS2 S-layer mixture was centrifugated at 23,000 xg for 40 minutes. The supernatant was discarded and the pellet containing S-layers was resuspended in fresh SEC buffer. This washing step was repeated five more times to ensure the removal of the detergent and contaminants. The purity of the sample was assessed using SDS-PAGE and ns-EM.

### Unfolding and refolding of the PS2 S-layer

Recombinant His-PS2^AD^ was purified as previously described. The gel-like fraction obtained after washing (described above) was resuspended overnight in unfolding buffer (8 M urea, 500 mM NaCl, 50 mM Hepes pH 7). The following day, the protein was loaded onto a SpinTrap column (Cytiva) pre-equilibrated with Buffer A, and washed five times with refolding buffer (500 mM NaCl, 50 mM Hepes, pH 7) with or without 10 mM EDTA. After washing, the proteins were eluted with Buffer B, also with or without 10 mM EDTA, and incubated overnight at 20°C before examination via ns- EM.

### Phylogenetic analysis

We assembled a database containing all 2325 Corynebacterium genomes and proteomes present at the GenBank database (Sayers et al., 2022) as of January 2024. We used HMM profile searches to identify protein PS2 in the protein database. First, we used the HMMER package (v3.3.2) (Johnson, Eddy, & Portugaly, 2010) tool jackhmmer to look for homologs of *C. glutamicum* PS2 in all the proteomes using the GenBank sequence AAX43986.1 as query. The hits were aligned with mafft (v7.475) (Katoh, Kuma, Toh, & Miyata, 2005) using default parameters. The alignments were manually curated, removing sequences that did not align globally. The hits obtained by jackhmmer might not include sequences that are very divergent from the single sequence query. For this reason, the alignment was used to create an HMM profile using the HMMER package (v3.3.2) tool hmmbuild. This specific and curated HMM profile of PS2 was used for a second and final round of searches against the proteomes using the HMMER tool hmmsearch. The new hits were aligned with linsi, the accurate option of mafft (v7.475), and trimmed using bmge (1.12) (Criscuolo & Gribaldo, 2010). The trimmed alignment was used to reconstruct the phylogeny of PS2. We repeated the search of PS2 against a database containing all *Corynebacteriales* order diversity (Gaday et al., 2022), obtaining no new hits. We inferred a maximum-likelihood tree of PS2 with IQ-TREE (Nguyen, Schmidt, von Haeseler, & Minh, 2015), using the posterior mean site frequency (PMSF) and the model LG + C60 + F + G, with ultrafast bootstrap supports calculated from 10,000 replicates. The guide tree required by the PMSF model was obtained using the LG+G+I+F model and the same trimmed alignment. To compare the genomic contexts of PS2, we retrieved 10 genes upstream and downstream of each PS2 hit, and we annotated the corresponding proteins using EggNOG-mapper (v2.1.12) (Huerta-Cepas et al., 2017) with the default parameters. The genomic context of PS2 in each strain was mapped on the *Corynebacterium* PS2 phylogeny using the online tool iTOL (Letunic & Bork, 2019) and custom scripts. We reconstructed a reference phylogeny of *Corynebacterium*, based on protein RNA polymerase subunit B, using the method described for protein PS2. We also reconstructed a reduced reference phylogeny of *Corynebacterium*, selecting only one strain per species (175 species), and a reduced phylogeny containing only the strains where PS2 was identified (102 taxa).

### Negative-stain transmission electron microscopy (TEM)

For visualisation of the PS2 S-layers by negative stain TEM, carbon-coated copper grids with 400-hole mesh (Electron Microscopy Science) were glow discharged (ELMO; Agar Scientific) with a plasma current of 5mA at vacuum for 60 s. Freshly glow-discharged grids were used immediately by applying 4 µl of sample (either purified PS2 or extracted directly from *C. glutamicum* cells) and allowing binding to the support film for 1 min after which the excess liquid was blotted away with Whatman grade 2 filter paper. The grids were then washed three times using three 15 µl drops of ddH2O followed by blotting of excess liquid. The washed grids were held in 15 µl drops of 2% uranyl acetate three times for, respectively, 10 s, 2 s, and 1min duration, with a blotting step in between each drop. Finally, the uranyl acetate-coated grids were fully blotted. The grids were then imaged using a 120 kV JEOL 1400 microscope equipped with LaB6 filament and TVIPS F416 CCD camera.

### Cryo-EM sample preparation and data collection

High-resolution cryo-EM dataset were collected using Quantifoil™ R2/1 300 copper mesh holey carbon grids. Grids were glow-discharged at 5 mA plasma current for 1 minute in an ELMO (Agar Scientific) glow-discharger. A Gatan CP3 cryo-plunger set at −176 °C and relative humidity of 90% was used to prepare the cryo-samples. Just before plunging, a DDM to a final concentration of 0.02 % was added to the ex-vivo purified PS2 S-layer solution and 3 μL was applied on the holey grid and incubated for 60 seconds. The sample was back-blotted using Whatman type 2 paper for 3 s and plunge-frozen into precooled liquid ethane at −176 °C. High-resolution movies were recorded at 300 kV on a JEOL Cryoarm300 microscope equipped with an in-column Ω energy filter (operated at slit width of 20 eV) automated with SerialEM 3.0.850. The movies were captured with a K3 direct electron detector run in counting mode at a magnification of 60K with a calibrated pixel size of 0.71 Å/pix, and exposure of 60e/Å2 taken over 60 frames. A total of 15208 movies were taken, of which 11988 were measured by tilting the stage at 30° and 3220 at 15° with a defocus range of -1.1 to -1.6 micrometers.

### Cryo-EM single particle analysis and structure determination

Movies were imported to CryoSPARC (Punjani, Rubinstein, Fleet, & Brubaker, 2017) where they were motion-corrected using Patch Motion Correction and defocus values were determined using Patch CTF. Exposures were curated and particles were picked using blob picker and extracted with a box size of 600 x 600 pixels. Several rounds of 2D classification were needed in order to clean selected particles providing a set of 1.268.481 high-quality particles and further centred at a hexameric axis. We performed non-uniform refinement using C6 symmetry. Next, we used two rounds of 3D classification in CryoSPARC and selected a single class showing higher-resolution information containing 521.917 particles. Finally, we used reference motion correction on those particles and the non-uniform refinement job to generate the final map with an average resolution of 2.44 Å according to the FSC curves (Supplementary Fig. 1). Finally, we used EMready (He, Li, & Huang, 2023) to improve the interpretability of the map in those regions where resolution was lower and finally, we built the atomic model using a combination of ModelAngelo (Jamali et al., 2024), AlphaFold2 (Jumper et al., 2021) followed by manually rebuilding in Coot (Emsley & Cowtan, 2004). A final round of refinement was performed using Phenix (Liebschner et al., 2019) and figures were done using ChimeraX (Pettersen et al., 2021). Map and model statistics are found in Supplementary Table 3.

### Phase contrast and fluorescence microscopy and image analysis

For imaging, a single colony of *C. glutamicum* ATCC13032 or strains expressing different variants of the PS2 S-layer were inoculated in 10 ml of LB media supplemented with 20 ml/L of Sodium lactate and with 25 µg/ml kanamycin when required and grown for 5h at 30°C. At this point, we added SpyCatcher-mCherry to a final concentration of 50 µM and the culture was grown overnight at 30°C with 120 rpm shaking. Next day, 10 ml of the overnight culture was harvested by centrifugation at 5000 x g for 5 minutes and washed 3 times with LB media to finally resuspend the pellet in 3 ml of fresh LB media. In a new culture tube, 2 ml of LB supplemented 20 ml/L of Sodium lactate and with 25 µg/ml kanamycin (when required) were inoculate with 500 µl of the above washed overnight culture. The culture was placed at 30°C for 1 hour after which SpyCatcher-GFP was added to a final concentration of 50 µM and incubated for 1h. For HADA labelling, cultures were incubated with 0.5 mM HADA for 20 min at 30 °C in the dark. Finally, the culture was harvested by centrifugation at 5000 x g for 5 minutes washed 3 times with 0.9% NaCl and diluted to OD600= 0.05 and 3 µL was transferred to the LB-agar strip. Images were collected in phase contrast and fluorescence mode on a Leica DMi8 inverted microscope (Leica) with 100X/1.32 oil objective (Leica). Phase contrast and fluorescent microscopy images were visualized and cropped using the software Fiji (Schindelin et al., 2012). They were segmented using the AI-based tool Omnipose (Cutler et al., 2022) specifically trained with a comprehensive dataset of *C. glutamicum* images, using the phase contrast channel. Masks were manually corrected and quantitative analyses were conducted with the Fiji plugin MicrobeJ (Ducret, Quardokus, & Brun, 2016) to generate fluorescent intensity heat maps and profile alignments. Heat maps represent the averaged localization of the fluorescent-tagged protein on a representative cell.

## Acknowledgements

We thank BECM and Dr. Marcus Fislage for his assistance during cryo-EM data collection. We thank Maxime Chazal for providing logistic support. A.S was supported by the EMBO (ALTF-709-2021) and the Marie Skłodowska-Curie Actions (MSCA; SLYDIV project). AW and JP were supported in part by grants from the Agence Nationale de la Recherche (ANR, France), contract ANR-21-CE11-0003, and Fondation pour la Recherche Médicale (FRM, contract EQU202303016284) and by institutional grants from the Institut Pasteur, the CNRS, and Université Paris Cité. Molecular graphics were rendered using UCSF ChimeraX, developed by the Resource for Biocomputing, Visualization, and Informatics at the University of California, San Francisco, with support from National Institutes of Health R01-GM129325 and the Office of Cyber Infrastructure and Computational Biology, National Institute of Allergy and Infectious Diseases.

## Author contributions

A.S., A.W. and H.R. designed the research. A.S. performed cloning and biochemical analysis. A.S. and M.S. performed NS-EM and Cryo-EM data acquisition and data analysis. A.S. acquired light microscopy images and J. P. performed data analysis. D. M. performed the phylogenetic and sequence analyses. A.S. and H.R. wrote the paper. All authors edited the paper.

## Data availability

The atomic coordinates and cryo-EM map have been deposited in the Protein Data Bank (PDB) and the Electron Microscopy Data Bank (EMBD) under the accession codes 9GK2 and 51414 respectively. Phylogenetic analysis data has been deposited to the Mendeley Data repository (doi: 10.17632/brj488xgky.1) and materials are available from the corresponding authors upon reasonable request. Requests for *C. glutamicum* strains should be addressed to the primary source, as cited in the manuscript.

## Competing interests

The authors declare no competing financial interests

## Notes

### Competing Interest Statement

The authors have declared no competing interest.

