## Supplementary Material for "Cryo-EM structure and polar assembly of the PS2 S-layer of *Corynebacterium glutamicum*"

### Affiliations:

### This file includes:

Supplementary Figures 1-10

Supplementary Table 1-3

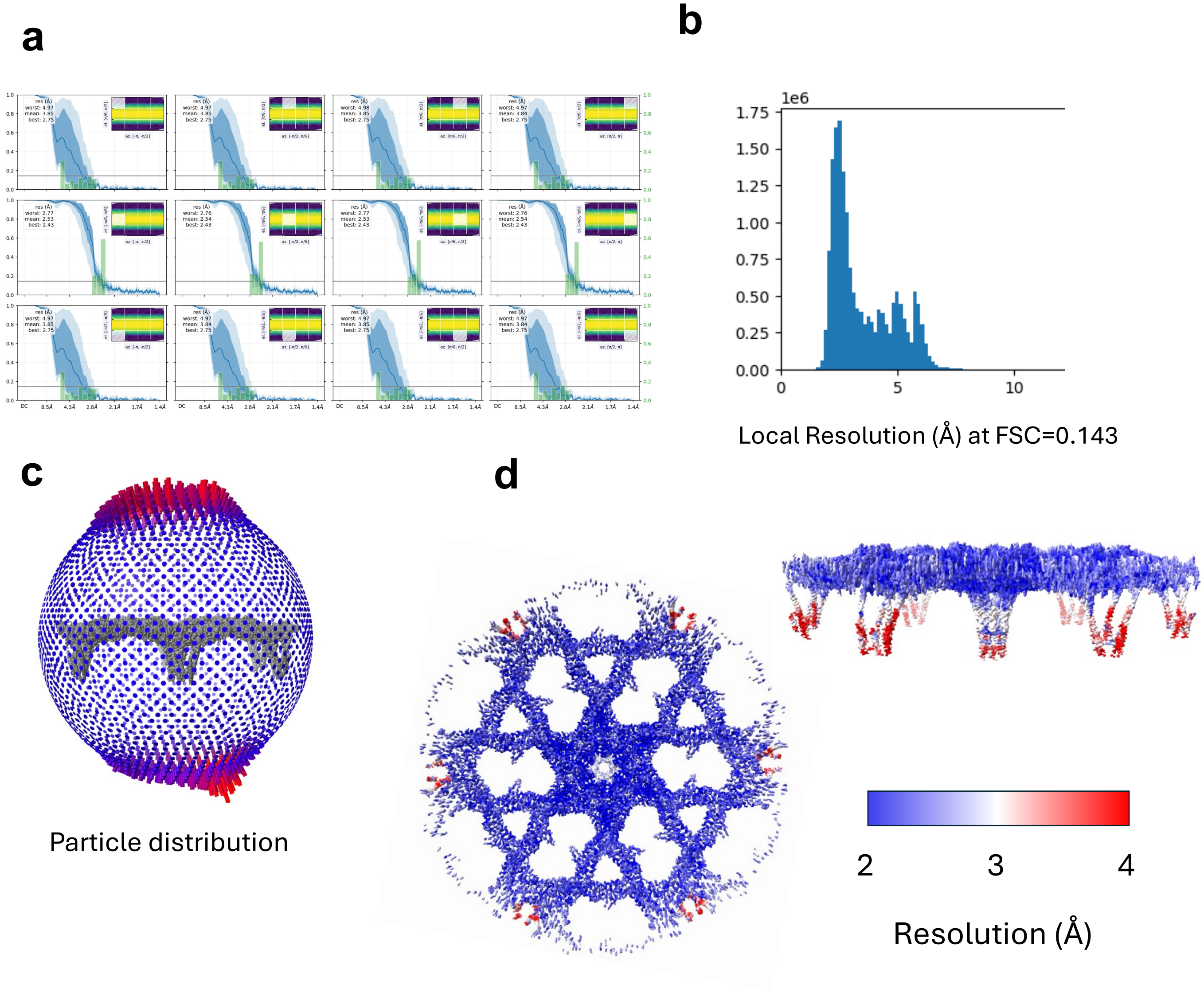

**Supplementary Figure 1. Single particle analysis of *ex-vivo* PS2 sheets. a.** FSC variation as a function of viewing direction obtained by the orientation diagnostics jobs (CryoSPARC) shows different map resolutions. **b**. Histogram of local resolutions in voxels of the cryo-EM map (C6-symmetrised). **c**. Particle distribution mapped on the cryo-EM density map. Red indicated hiher number of particles on that orientation, an enrichment of view at 15 and 30 degrees as a result of the tilted collection. **d**. Local resolution of the C2 cryo-EM map estimated in CryoSPARC, plotted into the density, shown from the top and side

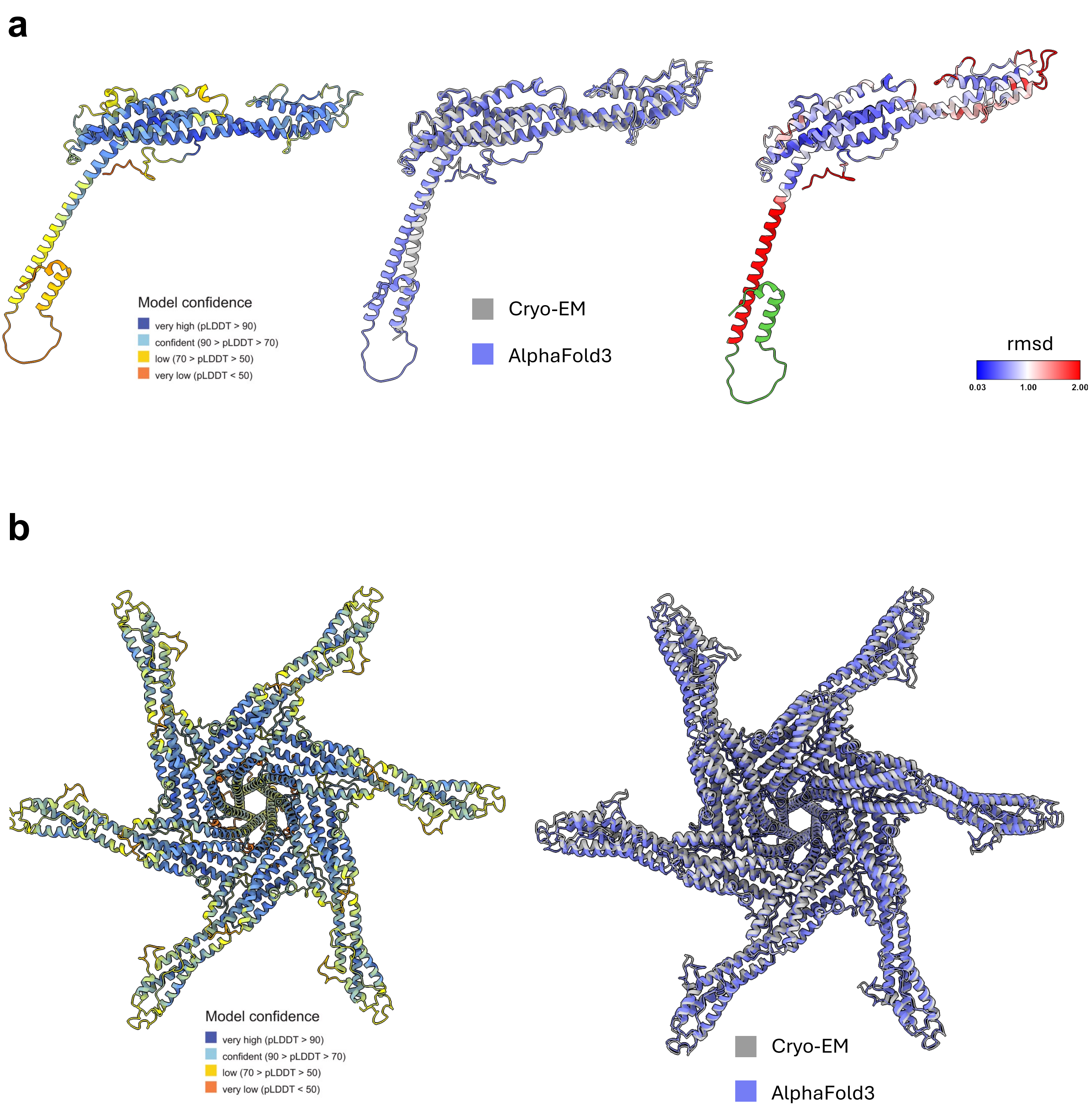

**Supplementary Figure 2. Prediction of the PS2 structure by AlphaFold3 compared with the Cryo-EM structure. a Left.** Ribbon representation of the PS2 monomer predicted by AlphaFold3, coloured by pLDDT values. **Middle.** Superposition of the predicted PS2 monomer (purple) with the Cryo-EM determined structure (grey) RMSD 3.28 Å. **Right.** AlphaFold3 model coloured by RMSD values when superimposed onto the Cryo-EM structure. Blue indicates close matches (low RMSD values), red indicates higher RMSD deviations, and green highlights areas lacking correspondence between the two structures. **b**. **Left**. Ribbon representation of the PS2 hexamer predicted by AF3 and coloured according to pLDDT values. **Right**. Structural superposition of the predicted AF3 structure with the experimentally obtained using cryo-EM. RMSD 2.76 Å.

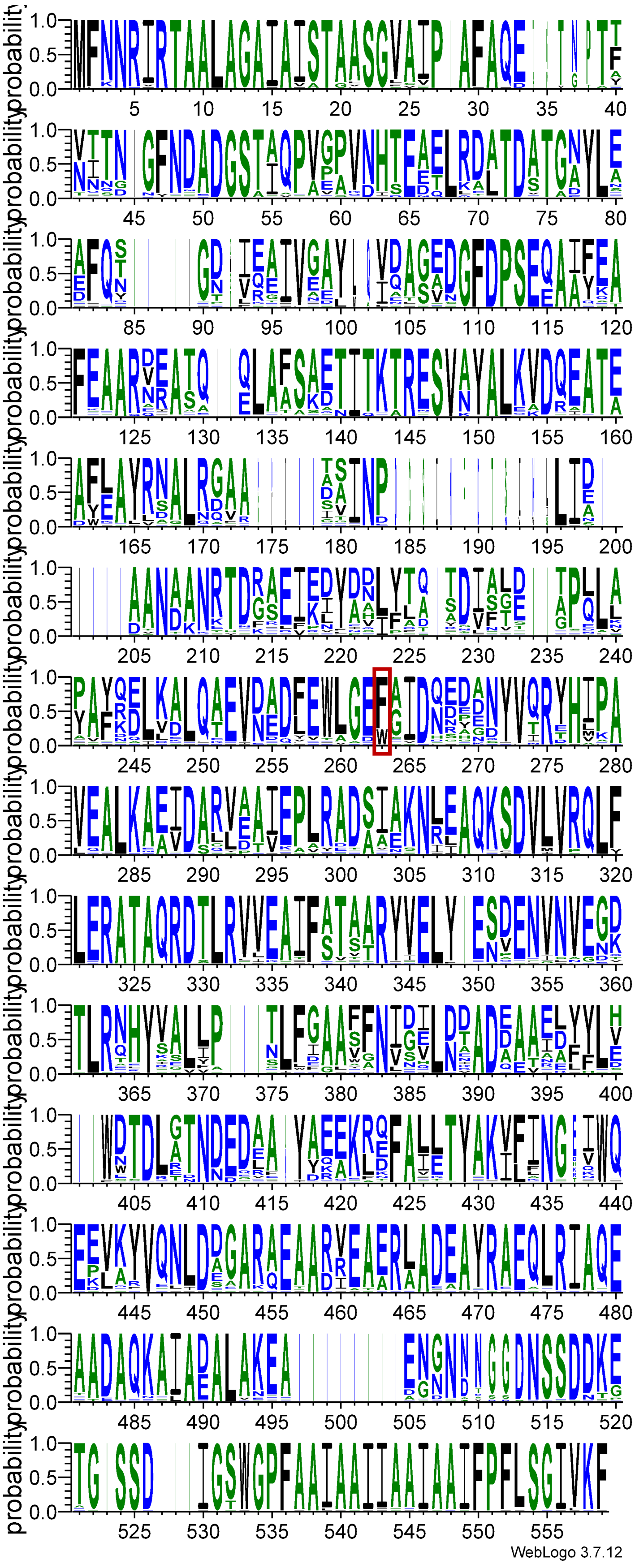

**Supplementary Figure 3. WebLogo of PS2.** Sequence alignment obtained from the ConSurf[1] server was used as input to generate the weblogo using WebLogo3[2]. Red boxed residue indicated the distal F from the arm region.

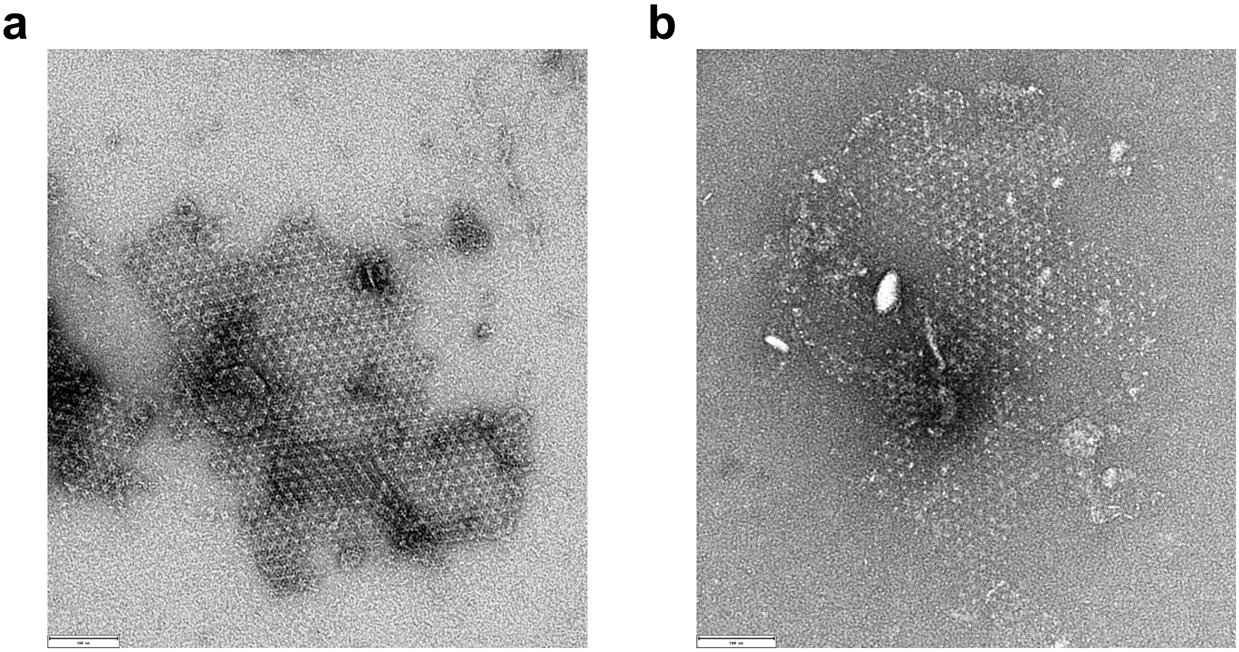

**Supplementary Figure 4. PS2$^{AD}$ S-layer assembly and stability are independent of divalent ions. a.** NS-EM of PS2 S-layer incubated with 10 mM EDTA for a week shows the presence of assembled S-layer. **b**. NS-EM of refolded PS2 in the presence of 10 mM EDTA. Scale bar 100 nm.

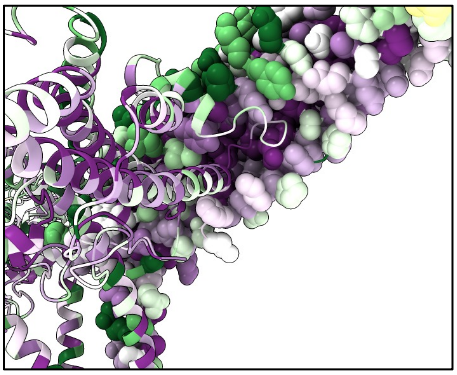

**Supplementary Figure 4. Conservation of the hexametric C6 interface.** The conservation of the PS2 S-layer has been mapped on the structure using the ConSurf server[1]. Purple indicated high levels of conservation. One PS2 promoter is shown in ribbon and the second one is shown as spheres.

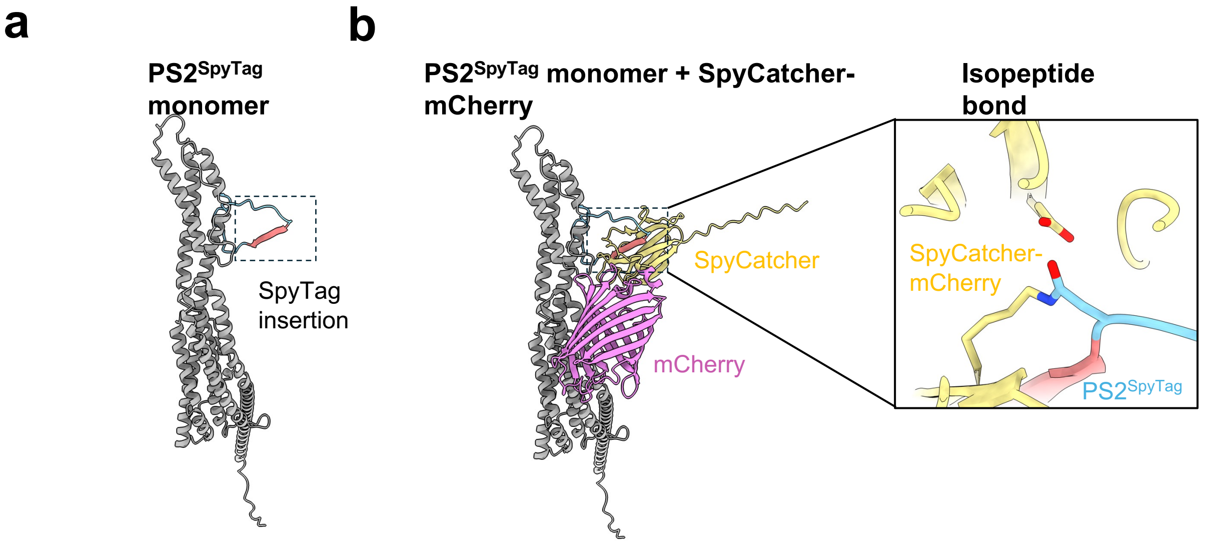

**Supplementary Figure 6. PS2$^{SpyTag}$ system. a.** AlphaFold model of the SpyTag added in the H3 insertion loop (left panel). **b.** Upon the exogenous addition of purified SpyCatcher:mCherry (middle panel) they form an isopeptide bond (covalent bond) catalyzed by a glutamic acid located in the proximity(right panel).

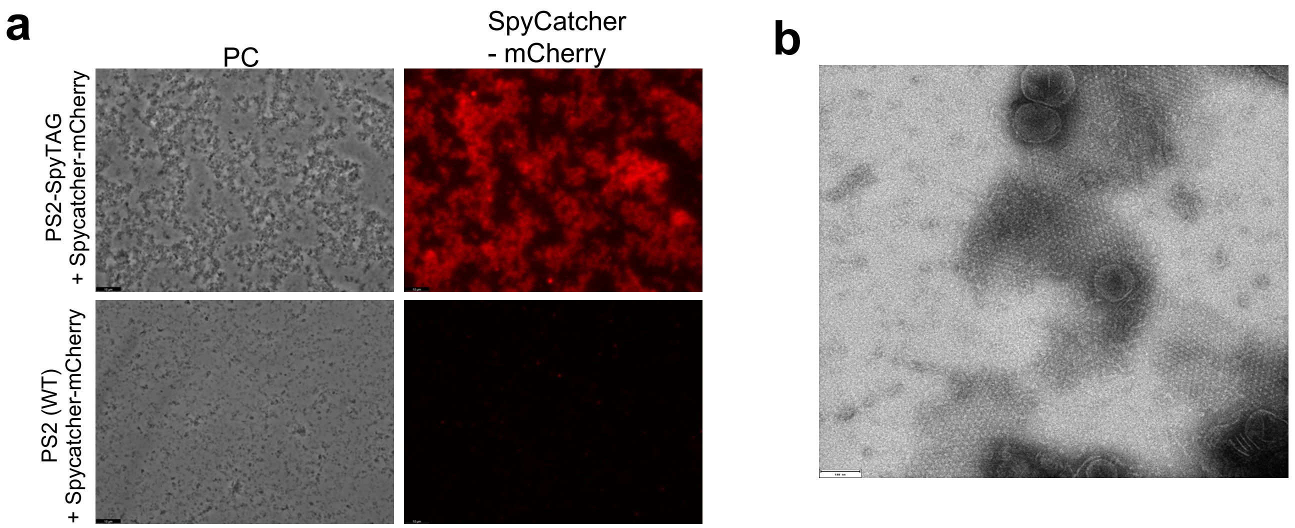

**Supplementary Figure 7.** ***In vitro*** **engineering of PS2. a.** Recombinant PS2-SpyTAG assembly domain is able to specifically bind SpyCatcher-mCherry (top) whereas the WT version shows no signal (bottom). Scale bar is 10 μM. **b**. PS2-SpyTag assembly domain still forms S-layer in vitro after being loaded with SpyCatcher-mCherry as observed using ns-EM. Scale bar is 100 nM.

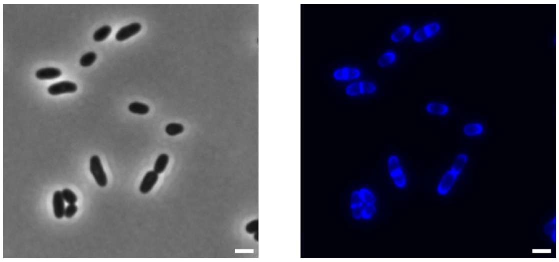

**Supplementary Figure 8. HADA labelling of** ***C. glutamicum*** **ATCC13032.** HADA is a fluorescent D-amino acid used to label newly inserted peptidoglycan. In *C. glutamicum*, peptidoglycan is synthesized at the poles and septum. Scale bar is 2 μM.

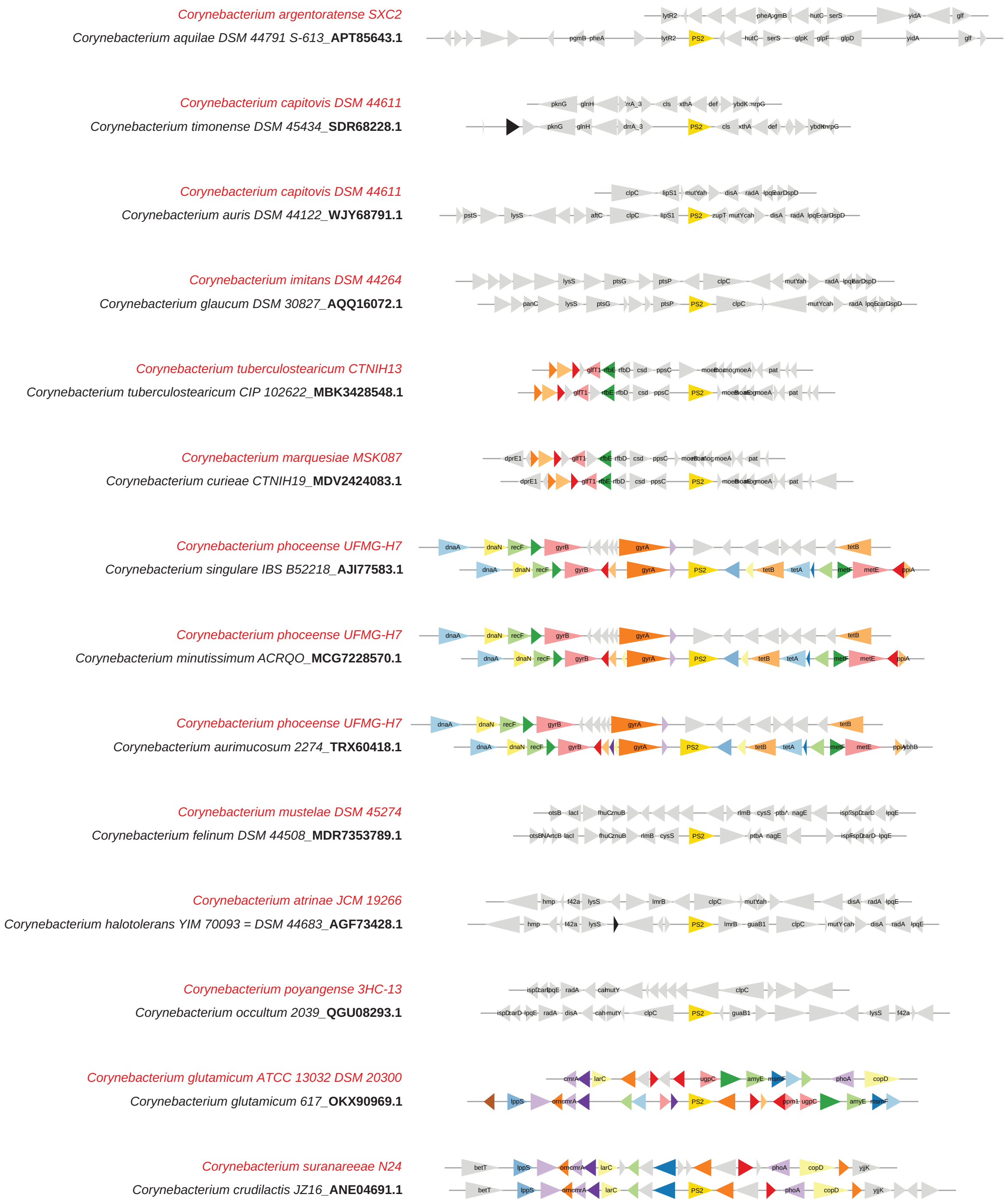

**Supplementary Figure 9.** Comparison of the genomic context of strains that code for PS2 and closely related strains that do not code for PS2. Triangles in colors correspond to genes frequently found in the same locus as *cspB* (PS2). Triangles without labels correspond to genes of unknown function.

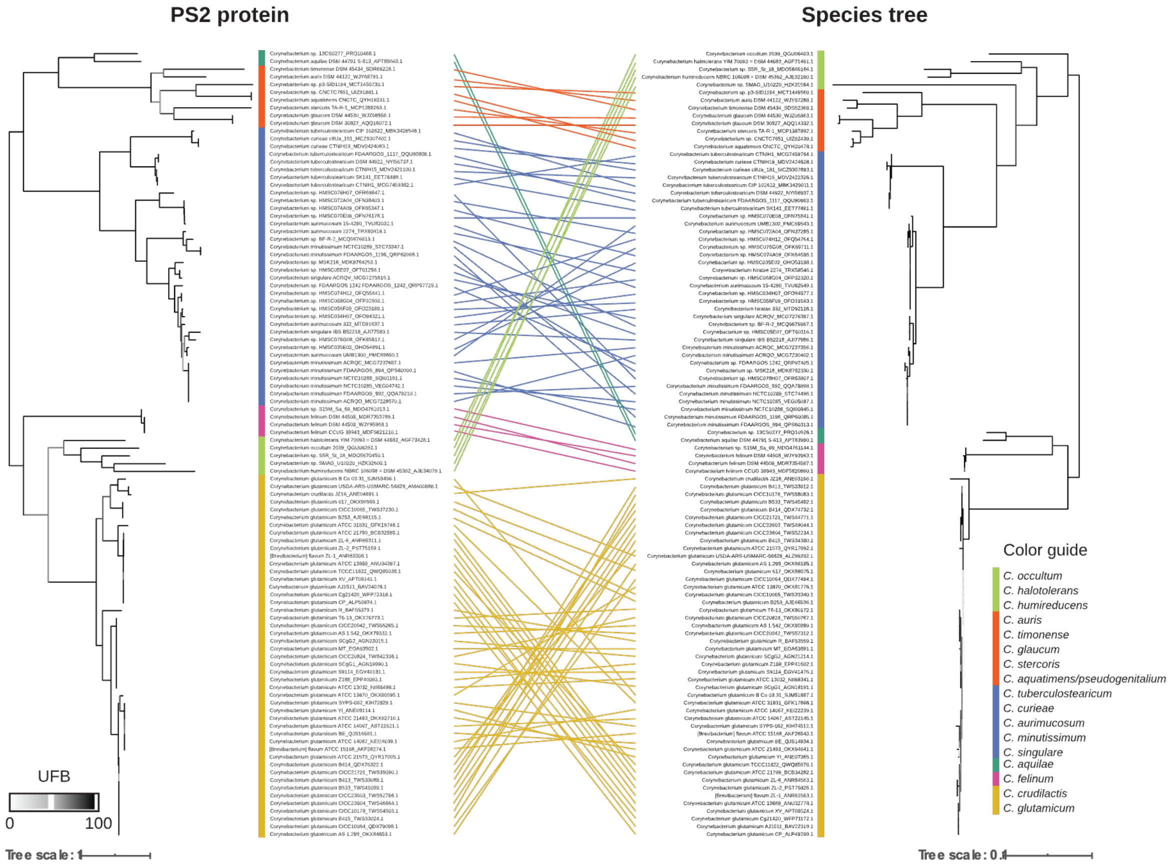

**Supplementary Figure 10.** Correspondence between the PS2 protein phylogeny and the species tree of the Corynebacterium genus. The species tree includes only the strains that contain protein PS2. The colors group species and strains that are monophyletic in the full Corynebacterium genus species phylogeny. The branches are colored in shades of gray according to the ultrafast bootstrap (UFB) supports.

| Oligo number | Name | Sequence (5' --> 3') |
|---|---|---|
| **p849** | F_PS2 | GGAGATATACAAATGCaggaaaccactgtaaccaccaacgg |
| **p868** | R_PS2_FL_V2 | CAGGTCAAGCTTAttaGTGATGATGATGGTGATGGAACTTAACGATACCGGAGAGGAATG |
| **p850** | R_PS2 | CAGGTCAAGCTTAttaGTGATGATGATGGTGATGCCAGGATCCGATGTCAGAAGAACCG |
| **p321** | F_open_pASK_3 | TAATAAGCTTGACCTGTGAAG |
| **p322** | R_open_pASK_3 | CATTTGTATATCTCCTTCTTAAAG |
| **p873** | F_Ps2_spyTAG_V3 | ATCGTCATGGTAGACGCCTACAAACGCTACAAAGGTTCTATCAACCCAGATACCTCTATCA ACCTACTGAT |
| **p874** | R_Ps2_spyTAG_V3 | CGTCTACCATGACGATGTGCGGCACGCCCCGACCGCCATCTGGGTTGATAGAGATAGC TGCATCG |
| **p862** | F_pTGR5_open | TAAAGCGGCCGCTTAAG |
| **p863** | R_pTGR5_open | AATATGCGGCCGCATATAT |

| Plasmid name | Vector family | Inducer | Resistance | Expressing gene | Reference |
|---|---|---|---|---|---|
| A231 | pASK-IBA3plus | anhydrotetracycline | Ampi | cspB assebly domain | This study |
| A232 | pASK-IBA3plus | anhydrotetracycline | Ampi | cspB full length | This study |
| A233 | pASK-IBA3plus | anhydrotetracycline | Ampi | spycatcher:sfGFP | This study |
| A234 | pASK-IBA3plus | anhydrotetracycline | Ampi | spycatcher:mCherry | This study |
| A235 | pASK-IBA3plus | anhydrotetracycline | Ampi | cspB assebly domain-SpyTag | This study |
| A236 | pTGR5 | N/A (cspB native promoter) | Kana | cspB full length | This study |
| A242 | pTGR5 | N/A (cspB native promoter) | Kana | cspB full length SpyTAG | This study |

| Strains | Characteristics | Source |
|---|---|---|
| ***E. coli*** | | |
| DH5$\alpha$ | F- endA1 Φ80dlacZΔM15 Δ(lacZYA-argF)U169 recA1 relA1 hsdR17(rK–mK+) deoR supE44 thi-1 gyrA96 phoA λ–; strain used for general cloning procedures | NEB (#C2987H) |
| BL21(DH3) | F- ompT hsdSB(rB–mB–) gal dcm (DE3); host for protein production | NEB (#C2527H) |
| ***C. glutamicum*** | | |
| ATCC13032 | Biotin-auxotrophic wild type | BCCM(#LMG 19741) |
| ATCC13032 *icd::cspB* | Biotin-auxotrophic wild type. Insertion of the cspB gene in the icd (isocitrate deshydrogenase) locus | Nicolas Bayan lab |
| ATCC13032 -PS2-SpyTAG | Biotin-auxotrophic wild type transformed with plasmid A242. Kana resistance. | This study |

**Supplementary Table 1.** The top table lists the oligonucleotides, the middle table lists the plasmids, and the bottom table lists the strains used in this study.

| Name | Protein Sequence |
| --- | --- |
| PS2 Full length | MFNNRIRTAALAGAIAISTAASGVAIPAFAQETNPTFNITNGFNDADGSTIQPVGPVNHTEETLRDLTDST<br>GAYLEEFQNGTVEEIVEAYLQVQASADGFDPSEQAAYEAFEAARVRASQELAASAETITKTRESVAYALK<br>VDQEATAAFEAYRNALRDAAISINPDGSINPDTSINLLIDAANAANRTDRAEIEDYAHLYTQTDIALETPQ<br>LAYAFQDLKALQAEVDADFEWLGEFGIDQEDGNYVQRYHLPAVEALKAEVDARVAAIEPLRADSIAKNL<br>EAQKSDVLVRQLFLERATAQRDTLRVVEAIFSTSARYVELYENVENVNVENKTLRQHYSALIPNLFIAAVA<br>NISELNAADAEAAAYYLHWDTDLATNDEDEAYYKAKLDFAIETYAKILFNGEVWQEPLAYVQNLDAGAR<br>QEAADREAARAADEAYRAEQLRIAQEAADAQKAIAEALAKEAEGNNDNSSDNTETGSSDIGSWGPFAA<br>IAAIIAAIAAIFPFLSGIVKF |
| PS2 Assembly domain | MQETNPTFNITNGFNDADGSTIQPVGPVNHTEETLRDLTDSTGAYLEEFQNGTVEEIVEAYLQVQASAD<br>GFDPSEQAAYEAFEAARVRASQELAASAETITKTRESVAYALKVDQEATAAFEAYRNALRDAAISINPDG<br>SINPDTSINLLIDAANAANRTDRAEIEDYAHLYTQTDIALETPQLAYAFQDLKALQAEVDADFEWLGEFGI<br>DQEDGNYVQRYHLPAVEALKAEVDARVAAIEPLRADSIAKNLEAQKSDVLVRQLFLERATAQRDTLRVV<br>EAIFSTSARYVELYENVENVNVENKTLRQHYSALIPNLFIAAVANISELNAADAEAAAYYLHWDTDLATN<br>DEDEAYYKAKLDFAIETYAKILFNGEVWQEPLAYVQNLDAGARQEAADREAARAADEAYRAEQLRIAQE<br>AADAQKAIAEALAKEAEGNNDNSSDNTETGSSDIGSW |
| PS2 signal peptide | MFNNRIRTAALAGAIAISTAASGVAIPAFAQ |
| PS2 SpyTag | MFNNRIRTAALAGAIAISTAASGVAIPAFAQETNPTFNITNGFNDADGSTIQPVGPVNHTEETLRDLTDST<br>GAYLEEFQNGTVEEIVEAYLQVQASADGFDPSEQAAYEAFEAARVRASQELAASAETITKTRESVAYALK<br>VDQEATAAFEAYRNALRDAAISINPDGGRGVPHIVMVDAYKRYKGSINPDTSINLLIDAANAANRTDRA<br>EIEDYAHLYTQTDIALETPQLAYAFQDLKALQAEVDADFEWLGEFGIDQEDGNYVQRYHLPAVEALKAE<br>VDARVAAIEPLRADSIAKNLEAQKSDVLVRQLFLERATAQRDTLRVVEAIFSTSARYVELYENVENVNVE<br>NKTLRQHYSALIPNLFIAAVANISELNAADAEAAAYYLHWDTDLATNDEDEAYYKAKLDFAIETYAKILFN<br>GEVWQEPLAYVQNLDAGARQEAADREAARAADEAYRAEQLRIAQEAADAQKAIAEALAKEAEGNNDN<br>SSDNTETGSSDIGSWGPFAAIAAIIAAIAAIFPFLSGIVKF |
| His6-SpyCatcher: GFP-StrepTag | MHHHHHHVTTLSGLSGEQGPSGDMTTEEDSATHIKFSKRDEDGRELAGATMELRDSSGKTISTWISD<br>GHVKDFYLYPGKYTFVETAAPDGYEVATPIEFTVNEDGQVTVDGEATEGDAHTSGGGGSSGSKGEELF<br>TGVVPILVELDGDVNGHKFSVRGEGEGDATNGKLTLKFICTTGKLPVPWPTLVTTLTYGVQCFSRYPDH<br>MKQHDFFKSAMPEGYVQERTISFKDDGTYKTRAEVKFEGDTLVNRIELKGIDFKEDGNILGHKLEYNFN<br>SHNVYITADKQKNGIKANFKIRHNVEDGSVQLADHYQQNTPIGDGPVLLPDNHYLSTQSVLSKDPNEK<br>RDHMVLLEFVTAAGITHGMDELYKGSWSHPQFEK |
| Name | DNA Sequence |
| PS2 promoter | GAATTCCTGTGAATTAGCCGGTTTAGTACTTTTCAGGGGTGTCTATTCTTACCAGATCGTCAAGTTGT<br>GGGTAGAGTCACCTGAATATTAATTGCACCGCACGGGTGATATATGCTTATTTGCTCAAGTAGTTCG<br>AGGTTAAGTGTATTTTAGGTGAACAAATTTCAGCTTCGGGTAGAAGACTTTCTATGCGCTTCAGAGCT<br>TCTATTAGGAAATCTGACACCACTTGATTAAATAGCCTACCCCCGAATTGGGGGATGGGTCATTTTTT<br>GCTGTGAAGGTAGTTTTGATGCATATGACCTGCGTTTATAAAGAAATGTAAACGTGATCAGATCGATA<br>TAAAAGAAACAGTTTGTACTCAGGTTTGAAGCATTTTCTCCGATTCGCCTGGCAAAAATCTCAATTGT<br>CGCTTACAGTTTTTCTCAACGACAGGCTGCTAAGCTGCTAGTTCGGTGGCCTAGTGAGTGGCGTTT<br>ACTTGGATAAAAGTAATCCCATGTCGTGATCAGCCATTTTGGGTTGTTTCCATAGCAATCCAAAGGTT<br>TCGTCTTTCGATACCTATTCAAGGAGCCTTCGCCTCT |

**Supplementary Table 2.** Sequences of interest used in this study. The SpyTag sequence is underlined.

| Appendix Table SX Cryo-EM model and data statistics | *Ex vivo* PS2 (EMDB-51414)(PDB 9GK2) |
|---|---|
| **Data collection and processing** | CryoARM300, BECM |
| Magnification | 60,000 |
| Voltage (kV) | 300 |
| Electron exposure (e–/Å$^2$) | 60 |
| Defocus range (μm) | -1.1 to -1.4 |
| Pixel size (Å) | 0.71 |
| Symmetry imposed | C6 |
| Tilts collected | 30 and 15 |
| Final particle images (no.) | 521.917 |
| Map resolution (Å) | 2.51* |
| FSC threshold | 0.143 |
| Map resolution range (Å) | 2.5-3.8 |
| **Refinement** | |
| Initial model used | AlphaFold2 |
| Model resolution (Å) | 2.44 |
| FSC threshold | 0.143 |
| Model resolution range (Å) | - |
| **Model composition** | |
| Non-hydrogen atoms | 60516 |
| Protein residues | 7776 |
| Ligands | 0 |
| **B factors (Å$^2$)** | |
| Protein (mean) | 53.80 |
| Ligand | NA |
| **R.m.s. deviations** | |
| Bond lengths (Å) | 0.004 |
| Bond angles (°) | 0.764 |
| **Validation** | |
| MolProbity score | 2.28 |
| Clashscore | 9.81 |
| Poor rotamers (%) | 4.10 |
| **Ramachandran plot** | |
| Favored (%) | 95.63 |
| Allowed (%) | 4.01 |
| outliers (%) | 0.36 |

* as from the 3d FSC job

**Supplementary Table 3.** Cryo-EM data collection, refinement and validation statistics of the PS2 structure.
